# *In vivo* characterisation of the toxicological properties of DPhP, one of the main degradation products of aryl phosphate esters

**DOI:** 10.1101/2019.12.26.888057

**Authors:** Samia Ruby, Jesús Marín-Sáez, Aurélie Fildier, Audrey Buleté, Myriam Abdallah, Jessica Garcia, Julie Deverchère, Loïc Spinner, Barbara Giroud, Sébastien Ibanez, Thierry Granjon, Claire Bardel, Alain Puisieux, Béatrice Fervers, Emmanuelle Vulliet, Léa Payen, Arnaud M. Vigneron

## Abstract

**Background:** Aryl phosphate esters (APEs), a main class of organophosphorus ester molecules, are widely used and commonly present in the environment. Health hazards associated with these compounds remain largely unknown and the effects of diphenyl phosphate (DPhP), one of their most frequent derivatives in human samples, are poorly characterised.

**Objective:** Our aim was to investigate whether DPhP *per se* may represent a more relevant marker of exposure to APEs and determine its potential deleterious biological effects in chronically exposed mice.

**Methods:** Conventional animals (FVB mice) were acutely (intravenous or oral gavage) or chronically (0.1 mg.mL^-1^, 1 mg.mL^-1^, 10 mg.mL^-1^ in drink water) exposed to relevant doses of DPhP or triphenyl phosphate (TPhP), one of its main precursors in the environment. Both molecules were measured in blood and other relevant tissues by liquid chromatography-mass spectrometry (LC-MS). Biological effects of chronic DPhP exposure were addressed through liver multi-omics analysis combining mRNA extraction and sequencing to high resolution LC-MS to determine the corresponding metabolic profile. Deep statistical exploration was performed to extract correlated information, guiding further physiological analyses (immunohistochemistry (IHC) and animal growth measurement).

**Results:** Acute and chronic exposure to DPhP led to significant levels of this molecule in blood and other tissues, an effect missing with TPhP. Multi-omics analysis confirmed the existence of biological effects of DPhP, even at a very low dose of 0.1 mg.mL^-1^ in drinking water. Chemical structural homology and pathway mapping demonstrated a clear reduction of the fatty-acid catabolic processes centred on acylcarnitine and mitochondrial β-oxidation. Interestingly, mRNA expression confirmed and extended these observations by demonstrating at all tested doses the overall repression of genes involved in lipid catabolic processes and regulated by PPARα, a master regulator of β-oxidation and its associated ketogenesis. IHC analysis confirmed the alteration of these pathways by showing a specific downregulation of Hmgcs2, a kernel target gene of PPARα, at all doses tested, and surprisingly, a strong reduction of the lipid droplet content only at the highest dose. Overall, DPhP absorption led to weight loss, which was significant using the highest dose.

**Conclusions:** Our results suggest that in mice, the effects of chronic exposure to DPhP, even at a low dose, are not negligible. Fatty acid metabolism in the liver in particular is essential for controlling fast and feast periods with adverse consequences on the overall physiology. Therefore, the impact of DPhP on circulating fat, cardiovascular and metabolic disease incidence deserves, in light of our results, further investigations.

## 1. Introduction

Di-phenyl phosphate (DPhP) has been used as a main biomarker for assessing exposure to aryl phosphate esters (APEs), especially tri-phenyl phosphate (TPhP), a molecule suspected of presenting human health hazards. However, this degradation compound can be produced from several APEs including ethylhexyl di-phenyl phosphate (EHDPhP) or the resorcinol bis(diphenyl phosphate) (RDP)^1, 2^ and tert-butylphenyl diphenyl phosphate (BPDP)^3^. Moreover, DPhP itself is largely present in the environment worldwide^4–8^, either owing to its spontaneous/microorganism production from known APEs^5, 9^, or to its direct use in industry^10^. Most APEs are used as flame retardants. They are added to consumer products and raw materials to delay combustion and meet flammability standards such as the ISO/TC92 Fire Safety, TC89 Fire Hazard existing in Europe. Moreover, unlike other flame retardants, TPhP and EHDPhP are also largely used as plasticizer and lubricants in hydraulic fluids, rubber, paints, textile coatings, food packaging and PVC, drastically increasing their presence in the environment. These compounds are not usually covalently linked to plastic materials and can easily leach into the environment^11^. High vapour pressure of TPhP is also likely to facilitate its release in the air once it is freed from its original material^12^. Not surprisingly, TPhP and EHDPhP are thus ubiquitous components of the human indoor environment where its sources and exposure pathways are quite diverse and heterogeneous with regards to other flame retardants. Indeed, TPhP and EHDPhP quantification in food, house dust, water or air has systematically demonstrated their presence in these very different matrices raising awareness on the safety of these compounds^4–8, 12, 13^. A study characterising the direct biological effects of DPhP and their relationship to TPhP exposure thus appeared to be of particular relevance to better define the effects and mechanisms of action associated with exposure to APEs in a more comprehensive way.

Historically, DPhP was believed to be produced from TPhP in the liver by oxidase and aryl esterase^14^. However, more recent analyses obtained from *in vitro* cultured hepatocytes, revealed that the main metabolites derived from TPhP were hydroxylated and glucuronated forms of TPhP^15^. Importantly, these metabolites and their equivalents for EHDPhP have recently been detected in human urine samples^16^, although DPhP remained the most abundant metabolite in these samples. This indicates either that degradation of APEs does not primarily occur in the human liver, or that APEs are rapidly degraded and absorbed as DPhP by the environment. Finally, the presence of DPhP in the environment could be directly due to its importance as a catalyst in polymerisation processes.

In line with these hypotheses, a recent study showed that serum hydrolase significantly contributes to TPhP and EHDPhP clearance and production of DPhP^17^. Bacteria present in the environment and the microbiota are likely to participate in this type of transformation since bacterial metabolism is able to fully degrade TPhP, DPhP initially being the main metabolite released in the biofilm^18^. Similarly, microsomal preparation of human skin demonstrated the ability of carboxylesterases to efficiently generate DPhP from TPhP^19^, indicating that likelihood of TPhP to reach subcutaneous fat and blood through this route of exposure is very low. Rapid detection of DPhP in urine samples of women exposed to TPhP through nail polish tends to confirm this hypothesis^20^. Finally, DPhP concentrations in the environment are strongly correlated with TPhP levels present in the same environment^4^, hence raising concerns about these potentially hazardous molecules for human health.

The complexity of the routes of exposure described above can cast doubts as to the relevance of *in vitro* and *in vivo* studies describing the toxicities associated with APEs such as the TPhP. For instance, very high doses of TPhP administered via oral gavage (300 mg/kg/day) in adult mice^21^ or through direct subcutaneous injection (around 200 µg/kg/day) in embryo/pups^22^ may artificially expose the organism to an irrelevant dose of TPhP and its hydroxylated forms, thus misrepresenting the more common route of DPhP exposure when APEs are present in the environment. These types of protocols have mainly led to the conclusion that TPhP has an obesogenic endocrine disrupting activity. These conclusions were reinforced *in vitro* by studies showing that high doses (10-100 µM) of TPhP can disturb the activity of the PPARγ^23, 24^, or by the ability of TPhP to enhance the lipogenic activity of the thyroid hormone on isolated chicken embryo hepatocytes^15^. Of note, these doses clearly show a high toxicity for mammalian cells casting strong doubts on the relevance of these results for human physiology. In addition, cell-based transactivation assays somehow failed to confirm agonistic or antagonistic activities of TPhP on either PPAR or TR nuclear receptors^25^. Moreover, a treatment combining 4 APEs administered at individual doses of 1 mg/kg/day, a protocol likely to expose animals to a relevant dose of DPhP, decreases the body weight gain of these animals instead of increasing it. Similarly, recent reports indicated that exposure DPhP and TPhP could disrupt the metabolism in opposite manners^22^.

On these bases, we estimated that a large study mimicking optimal and relevant routes and doses of DPhP exposure in mouse models was critical. To validate our choice of using DPhP rather than TPhP or another APE in our toxicity study, we first analysed DPhP concentrations in blood of mice treated with various doses of both molecules via different routes of exposure. We hypothesised that humans are more likely to be continuously/chronically exposed to TPhP and DPhP owing to the presence of TPhP in air and dust, rather than temporarily/acutely exposed through nutrition. We thus decided to analyse how these two molecules were absorbed more continuously though drink water and kinetically transformed in mice, in comparison with other acute modes of administration such as oral gavage or tail-vein injection. We then presented the data reporting the bioaccumulation and distribution of these molecules in mice. Finally, since our aim was to analyse the biological consequences of a relevant DPhP exposure, we defined a workflow based on multi-omics analyses combining metabolomics and transcriptomic analyses on tissue extracts obtained from independent experiments, followed by a histological validation. Results clearly demonstrate the ability of DPhP at a very low dose to disturb lipid metabolism processes in the liver, strongly questioning the safety of APEs.

## 2. Results

### Correlation analysis between TPhP/DPhP exposure and their level in blood and liver

When 0.1 or 1 µg TPhP was injected directly into vein-tail or administered by oral gavage, DPhP was detected after one hour in a dose-dependent manner in whole blood (Figure 1A, B). Inversely, at these concentrations, TPhP could not be detected in the blood of exposed animals (data not shown). After administration of 10 µg or 100 µg TPhP, TPhP was only quantified in the blood of two animals at 2.33 ng/mL and 10.20 ng/mL, exposed to 100 µg following intravenous injection and oral gavage, respectively. In all other animals (18/20 animals), TPhP remained undetected. These results indicate that TPhP was either, rapidly transformed in the bloodstream or in the gut by the microbiota, or not absorbed by the digestive tract. To determine whether TPhP transformation into DPhP was the reason for the lack of detection of TPhP in the bloodstream, we also quantified DPhP in these same experiments. We detected small quantities of DPhP with the highest dose of TPhP administered (Figure 1A, B), but these were negligible compared to concentrations obtained after direct DPhP exposure, suggesting that detection of DPhP is not generally the consequence of the transformation of TPhP.

**Figure 1.**
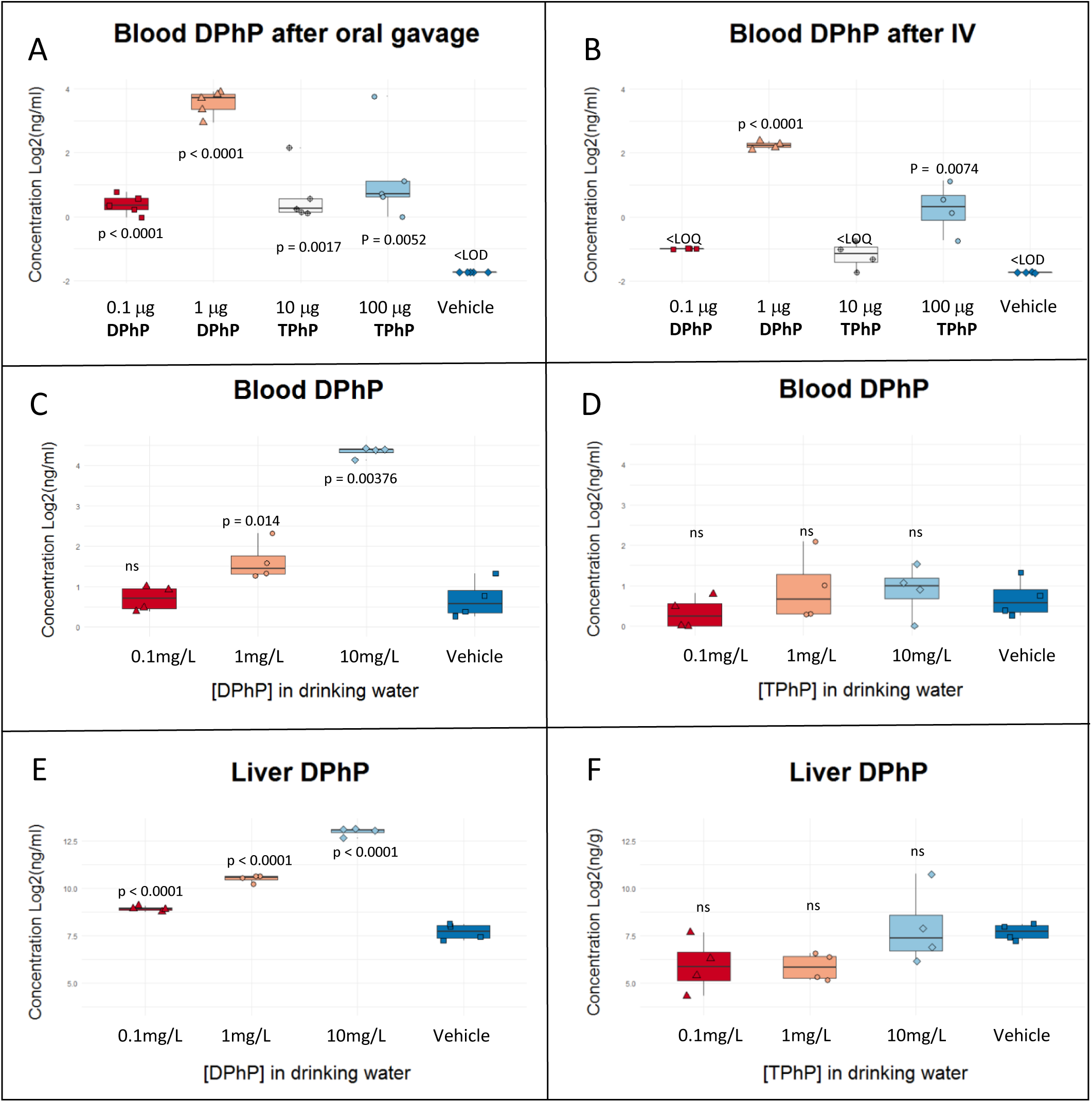
Acute exposure to DPhP and TPhP A-D. 5 animals were exposed to the indicated doses of DPhP or TPhP via two routes of administration. After 1 h, DPhP was quantified from the whole blood. Concentrations obtained are plotted on the box and whisker plot indicating significant p-value calculated between tested conditions and the vehicle. E-F. Identical experiment with DPhP quantification performed on liver extracts.

We further compared these results to a more continuous exposure to both molecules present in the drinking water of animals to mimic their chronic ingestion from swallowed dust. We used 3 concentrations of 0.1 mg/L, 1 mg/L and 10 mg/L, equivalent to 0.5 µg, 5 µg and 50 µg of each molecule ingested overnight (active period for mice). These quantities were comparable to those used with other routes of exposure. At the two highest concentrations, DPhP was still dose-dependently detectable in whole blood with lower concentrations measured than those obtained previously (Figure 1C). This was expected since the dose was now spread over a much longer period. Inversely, we did not detect any significant amount of TPhP in animals exposed to this same molecule through drinking water (data not shown). Most importantly, DPhP was not retrieved from the whole blood of animals exposed by this route to TPhP (Figure 1D).

Since the liver is the first organ to process exogenous molecules absorbed from the digestive tract, we further determined if these molecules were present in this organ using the same exposure doses via drinking water. DPhP was detected and quantified in a dose-dependent manner at all concentrations tested (Figure 1E), indicating that the molecule is efficiently absorbed in the liver before its eventual bio-transformation and clearance by the kidney. Again, neither TPhP nor DPhP were detected in the liver of animals exposed to TPhP through drinking water (Figure 1F), confirming that TPhP ingestion is not at the origin of the DPhP found in human urine.

### Bio-accumulation and distribution of DPhP in chronically exposed mice

Based on these analyses, we next focused on the consequences of chronic exposure to DPhP. Mice were exposed daily to similar doses of DPhP in their drinking water, over a 12-week period. We analysed four tissues, namely the whole blood, the liver, the visceral fat and the mammary gland, potentially presenting a tropism for the molecule, either due to their chemical characteristics (presence of hydrophobic constituent for mammary gland and visceral fat) or to the route of exposure used for these experiments (whole blood and liver). In the blood, DPhP was again significantly detected in all animals chronically exposed to DPhP in drinking water at 1 mg/mL and 10 mg/mL. (Figure 2A). Values obtained after chronic treatment were significantly higher in comparison with our previous overnight exposure (Figure 1E and 2A). For instance, exposure to 1 mg/mL resulted in an increase (> 2-fold), indicating that a cumulative effect occurred during chronic exposure. DPhP was also detected in the liver at all tested concentrations in a dose-dependent manner (Figure 2B). However, a cumulative effect was not observed here, suggesting that other tissues had likely absorbed the molecule and/or released it into the bloodstream. Consistently, DPhP was detected in both visceral fat and mammary gland, gradually increasing with treatment doses (Figure 2C, D).

**Figure 2.**
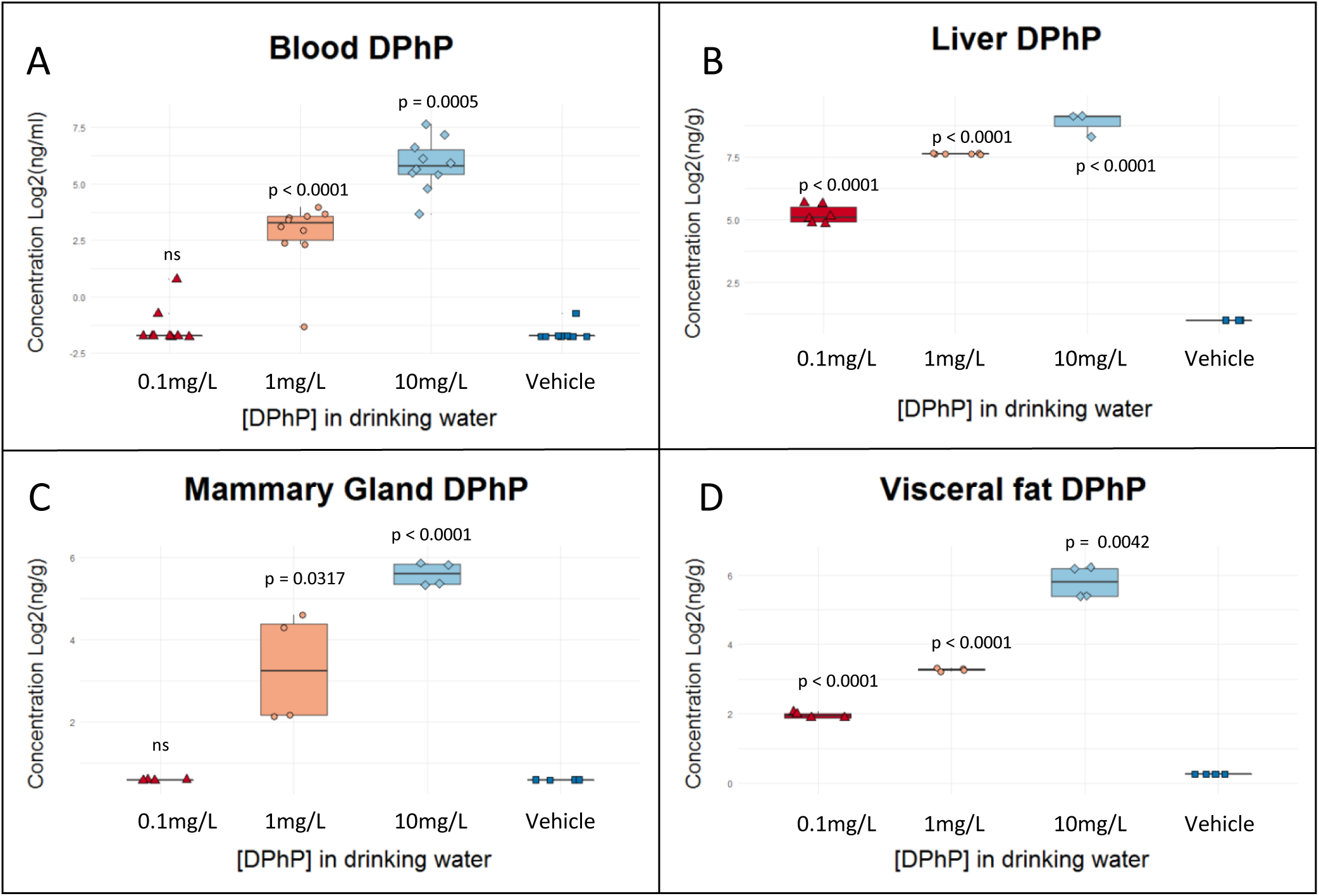
Chronic exposure to DPhP A-D. 5 animals were chronically exposed to the indicated concentrations of DPhP for 12 weeks through drinking water. DPhP was then quantified in the indicated biological matrix. Concentrations obtained are plotted on the box and whisker plot indicating significant p-values calculated between tested conditions and the vehicle.

### Biological effects of exposure to DPhP through metabolomics analyses of the liver

Since DPhP was abundant in the liver of all treated animals, even at the lowest concentration of 0.1 mg/L given via drinking water, we focused our subsequent experiments on this organ in order to measure the possible biological consequences of these molecules. Through multiple injections (n = 8), we first verified that the metabolic profiles obtained by LC-HRMS could discriminate the 4 groups of samples (untreated (CTRL) or treated with 3 different doses of DPhP, 0.1 mg/L (C1), 1 mg/L (C2), 10 mg/L. (C3)). Principal component analysis applied using the unit variance algorithm discriminated the samples, 68% of the variance being attributable to the first 3 axes of the PCA (Figure 3A). To determine which metabolites were involved in this discrimination, we used a volcano plot representation to compare each treated group to the control animals and annotated the most discriminating compounds as indicated in the Figures 3B-D. At this stage, discriminating compounds were identified through their exact mass, their expected presence in mammalian organisms and, eventually their fragmentation spectrum (see methods and Figure S1). As expected, discriminating compounds were relatively heterogeneous, chemically and functionally, even though a subset of acylcarnitines were apparently significantly downregulated in exposed animals at all concentrations. From these first hits, we decided to perform an in-depth analysis combining 175 annotated metabolites either highly enriched or showing a very robust deviation from control conditions, to which we added the main intermediate metabolites (glycolysis, TCA cycle, amino-acids, fatty acids…) (Supplemental Table 1). Due to the heterogeneity of these metabolites and, for some of them, their non-appurtenance to the endogenous metabolism, we performed an initial comparison based on their structural homology and chemical ontology. This type of analysis eliminates bias due to incomplete mapping and network size heterogeneity observed when using classical pathway enrichment^26^. A circular tree plot for each concentration is shown in Figure 4A. One of the clusters clearly highlighted acylcarnitines, an important amount of these metabolites showing a clear trend towards a lower concentration after exposure to DPhP. Interestingly, another pool of increasing metabolites appeared in a dose-dependent manner, revolving around the purine nucleotides and containing metabolites associated with the synthesis of nicotinamide dinucleotide. To extend these analyses, we then constructed an enrichment plot displaying the significance of the observed change for structural homologies, the size of the cluster and the homogeneity of their variation (Figure 4B and S2A, B). We confirmed a strong and significant decrease in acylcarnitines for all DPhP concentrations. In line with these results, an important number of fatty acids were altered, even though the direction of these alterations was not homogeneous. Moreover, we also noted an increase in purine-relative metabolites, even for the lowest DPhP concentration with a significant dose-dependent increase (cluster purine, purinone, nicotinic acid). Next, to determine how these different clusters were organised around endogenous metabolic networks, we combined chemical and biochemical mapping in a joint analysis using the MetaMapp algorithm^27^ (Figure 4C and S2C, D). A clear connection was observed between the reduction of acylcarnitine pool and the decrease in a subset of fatty acids. Conversely, the increase in purine and dinucleotide metabolism was more scattered in the network mapping nucleic base, and tryptophan metabolism clusters. Moreover, we noted the existence of a metabolite pool containing endogenous and exogenous molecules with aromatic cycles known to be controlled by xenobiotic-activated responses. We then decided to perform a hierarchical clustering of our annotated metabolites associated with their detected level in each condition, the aim being to combine on the same graph the results obtained with the different concentrations of DPhP used. Interestingly, this analysis performed with only 175 metabolites clearly discriminated the 3 exposed conditions from the control, in a similar way to the PCA based on the entire dataset of detected mass (Figure 3A and 4D). Moreover, the metabolic profiles described in this analysis and associated with exposed animals strongly clustered together in comparison with the control, with the lowest concentration being the least distant. The intermediate and the highest concentrations gave the closest results, but a careful inspection of the obtained clusters confirmed the reduction of the acylcarnitine pool even with the lowest concentration of DPhP and without a clear dose-dependent type response (Figure 4E). The most abundant fatty acids such as oleic acid and palmitic acids were also present in this cluster. Inversely, another cluster containing various types of molecules clearly displayed metabolite accumulation in a dose-dependent response (Figure 4F). Several xenobiotics were present in this cluster, as well as tryptophan and aromatic derivatives indicating that exposure to DPhP, itself a xenobiotic and an aromatic compound, was disturbing the metabolism associated with these molecules. Finally, we noted the presence in this cluster of dodecanedioic acid. The accumulation of this metabolite is associated with an impairment of fatty acid oxidation at the carnitine palmitoyl transferase (CPT) level^28^, an effect in line with the observed overall reduction of acylcarnitine pools.

**Figure 3.**
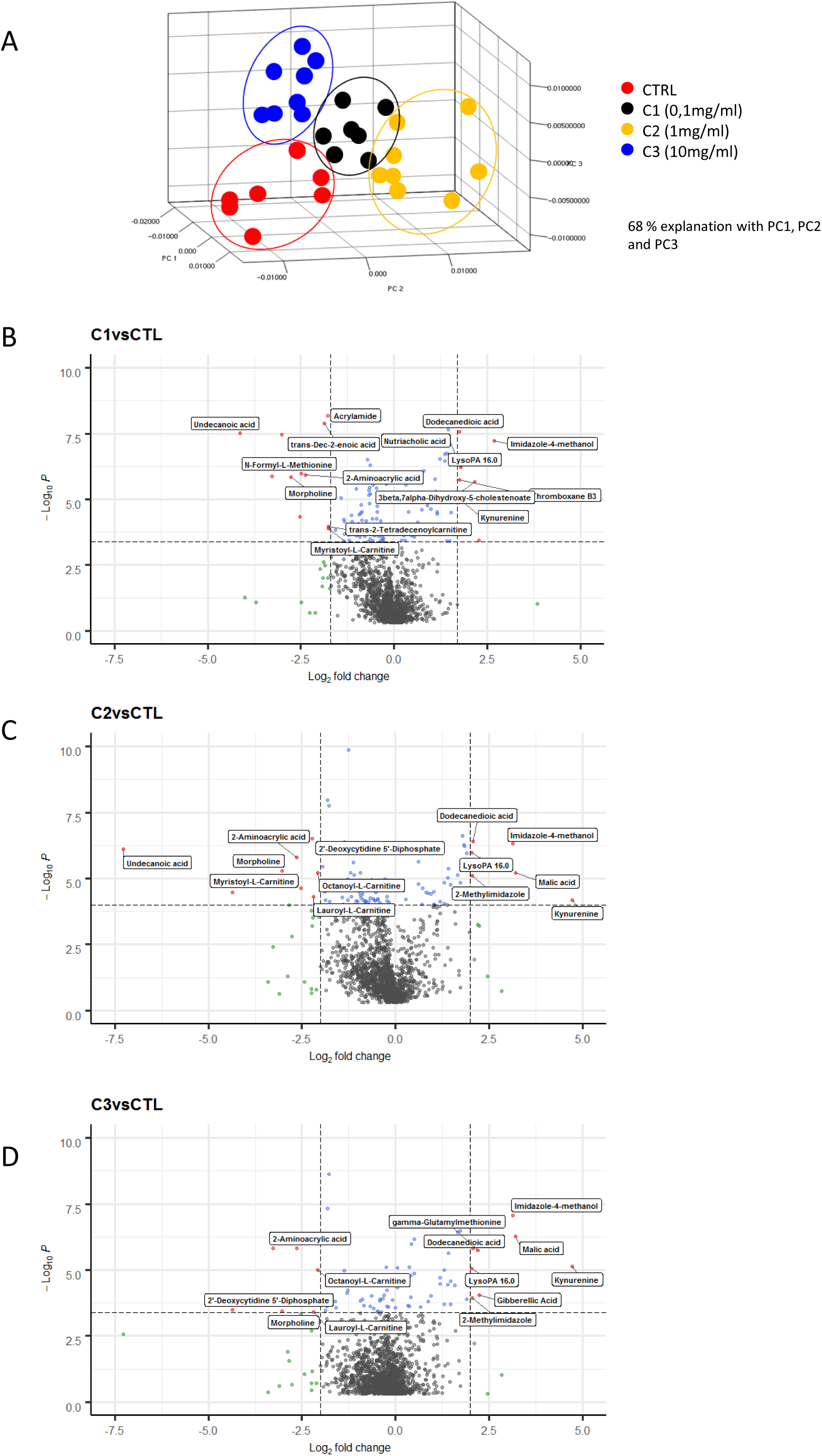
Hepatic metabolomics analysis of animals exposed to DPhP A. Principal Component Analysis was performed using pareto algorithm as an observatory method to discriminate groups of samples based on the amount of retained ion m/z (see methods). Explained variance with the 3 first axes is indicated. B-D. A volcano plot was created through supervised partial least squares (PLS) (using pareto algorithm) and T-Test (using group mean algorithm), individually comparing the metabolite concentrations for each dose of DPhP used against control values. Most significant metabolites were annotated when possible and indicated directly on the plot.

**Figure 4.**
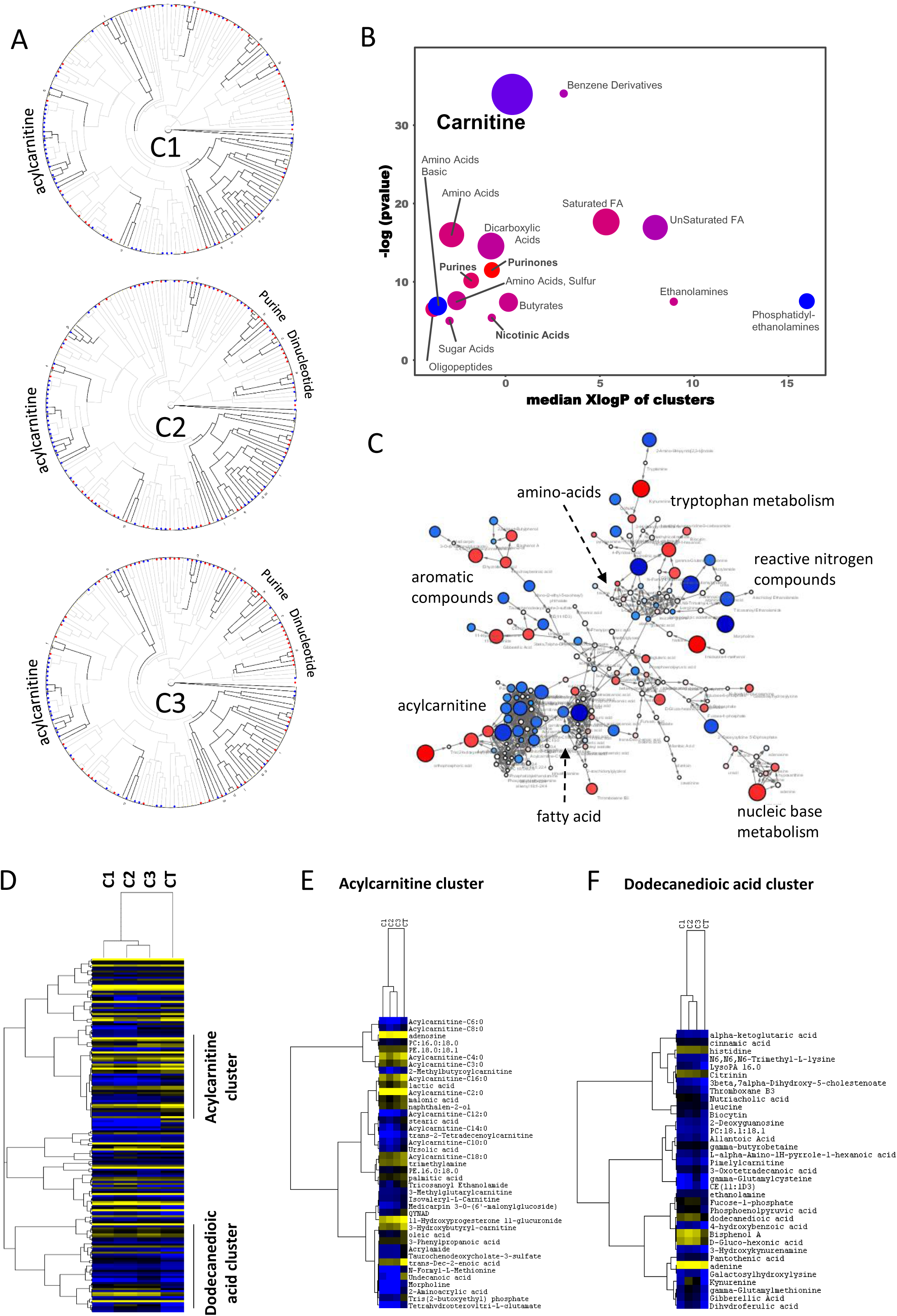
Metabolic network analysis through structural identity and pathway mapping A. Data associated with 175 metabolites either significantly altered or belonging to the main intermediate metabolites were used to perform a Chemical Similarity Enrichment Analysis (ChemRICH). A Tanimoto chemical similarity mapping form a clustered circular similarity tree. Dark black lines indicate boundaries of clusters that are significantly different in exposed animals at the indicated concentration of DPhP versus control mice (p < 0.05). Increased metabolite levels in exposed mice are labelled as red nodes, decreased levels are marked in blue. Cluster label is indicated. B. ChemRICH set enrichment statistics plot for the same metabolites as A, extracted from exposed animals with the C2 concentration of DPhP versus control. Each node reflects a significantly altered cluster of metabolites. Enrichment p-values are given by the Kolmogorov– Smirnov-test. Node sizes represent the total number of metabolites in each set of clusters. The node colour scale shows the proportion of increased (red) or decreased (blue) compounds in exposed mice compared to control mice. Purple-colour nodes have both increased and decreased metabolites. C. MetaMapp visualization of the same metabolomics data highlighting the differential metabolic regulation and the organization of metabolic clusters based on KEGG reactant pair information and Tanimoto chemical similarity matrix. Increased metabolite levels in exposed mice are labelled as red nodes, decreased levels are marked in blue. Intensity differences are also encoded in node size. Cluster label is indicated. Metabolites previously clustered together based on their structural similarity could now be separated according to their different pathway mapping (dinucleotides are now present in the tryptophan metabolism cluster, whereas purine are in the nuclei base metabolism.) D-F. Identical data obtained for the same 175 metabolites and the same 4 groups of exposed and control animals were clustered hierarchically through their relative level (complete linkage with spearman correlation). Yellow-blue encoding is used to represent these metabolites according to their absolute amounts. Distance between levels of samples and metabolites are shown as two tree plots. Clusters containing the acylcarnitines or the dodecanedioic acids are highlighted and enlarged in the indicated right-hand panels.

Overall, these results indicate that fatty acid oxidation is disturbed upon exposure to DPhP, combining a lack of fatty acid substrate such as oleate, and a reduction of enzymatic activities required for acylcarnitine production from these substrates. Moreover, as shown above, increasing doses of DPhP apparently disturbed the xenobiotic response and the metabolic connections between the purine, the dinucleotide and the tryptophan metabolism. In order to validate these conclusions, we performed a last series of analyses to confirm the most important metabolites associated with this study. Retention time and fragmentation spectra of standards were obtained and compared to the values obtained from the samples. Supplemental Table 2 recapitulates the confirmed metabolites and their mean and median concentrations obtained with the different animals exposed.

### Biological effects of DPhP exposure through transcriptomic analyses of the liver

The second part of our multi-omics analyses was then performed on another batch of 20 animals treated identically. mRNA were extracted from pooled liver and analysed by next-generation sequencing using 4 analytical replicates. Genes were then filtered and only those displaying a mean RPKM > 0.2 in the control condition were conserved (Supplemental Table 3). PCA was performed using the mean expression associated with the 4 experimental conditions. When the first axis was considered, the three treated conditions were significantly different from the control, the most discriminating treatment being associated with the intermediate concentration of DPhP (Figure 5A left panel). Interestingly, this pattern was correlated with the metabolic difference observed, since this concentration had the greatest effects. We noticed that 44% of the total variance was attributable to the first axis (Figure 5A right panel), suggesting that the genes with the highest Eigen values on this axis were the most relevant for the biological effects of DPhP. We consequently selected the most discriminating genes (Supplemental Table 4, Eigen value > 0.3) and performed a gene ontology analysis using the String software v11. For the analysis, genes were ranked according to their Eigen value for generating a functional enrichment score/false discovery rate, then used to construct a volcano plot (Figure 5B and Supplemental Table 5). The most significantly and highly depleted processes in the exposed versus control conditions, were those related to lipid metabolism and more specifically to fatty acid oxidation. However, we could also notice a significant inhibition of genetic response associated with xenobiotic metabolism. These results were thus indicative of the existence of an inhibition of these genetic programmes in exposed animals and of a relevant correlation with our previous experiments analysing the metabolome of animals treated identically. Moreover, when we retrieved the gene list associated with the fatty acid catabolic process, and selected the most discriminating genes in our dataset (Eigen value > 0.3), we confirmed that these genes encoded functional protein networks related to mitochondrial and peroxysomal fatty acid oxidation with a very high confidence rate (Figure 5C left panel). Since fatty acid oxidation is strongly regulated by the PPAR transcription factor, we verified that PPAR signalling was also among the significant terms associated with our ranked gene list (Figure 5C right panel). The network reconstructed from this last term demonstrated that PPARα target genes were at the heart of the dysregulation process, strongly suggesting a specific alteration of this genetic response critical for the control of lipid catabolism in the liver.

**Figure 5.**
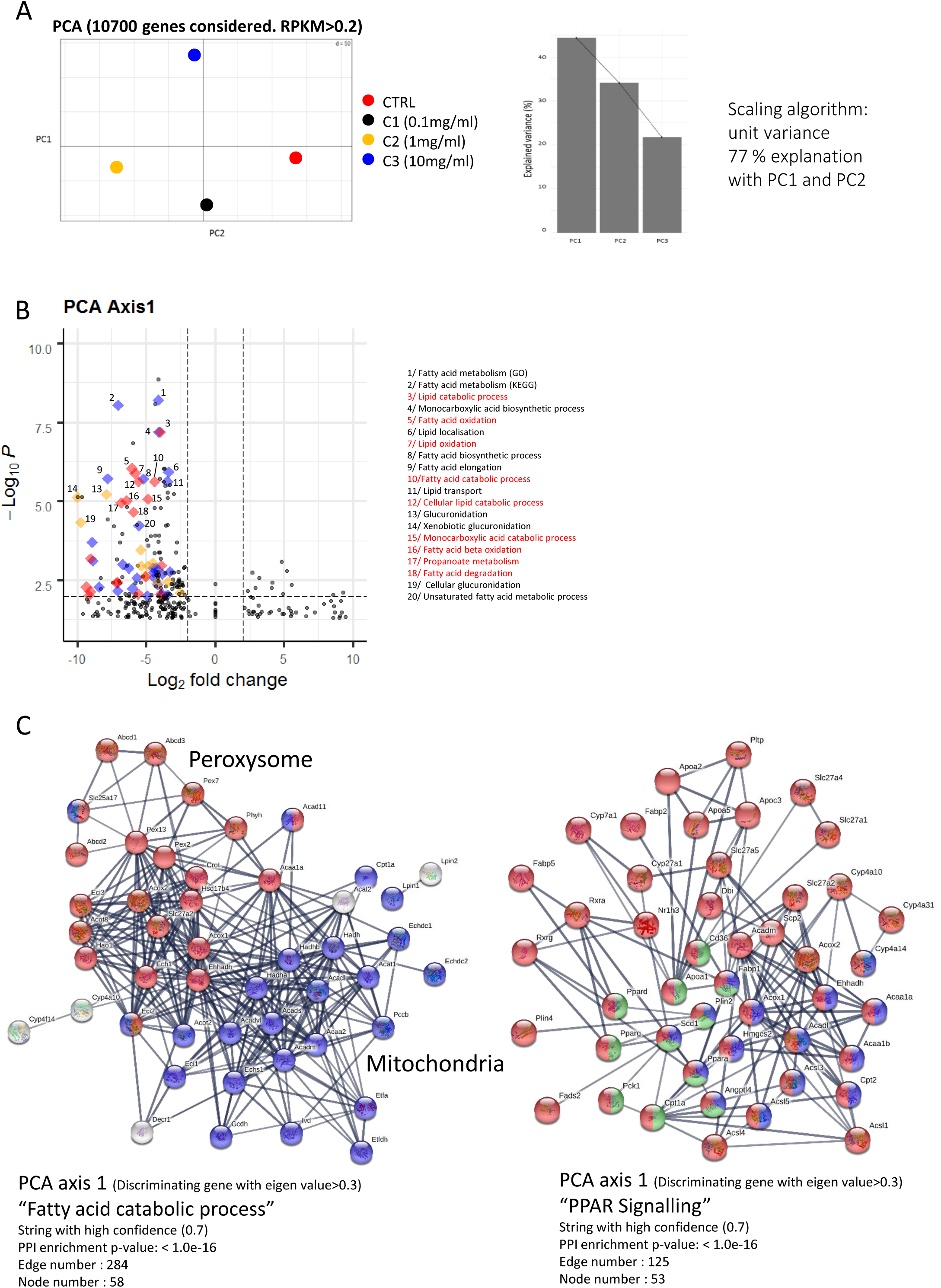
Transcriptomic analysis of liver belonging to animals exposed to DPhP A. Principal Component Analysis was performed using pareto algorithm as an observatory method to discriminate group of sample replicates based on the amount of mRNA expression obtained from indicated animals, through reverse transcription and Next-Generation- Sequencing. Considered genes and explained variance with the 2 first axes of the PCA are indicated. B. Gene ontology analysis of the most discriminant genes (Eigen value > 0.3) used to build the first axis of the previous PCA performed through Functional Enrichment Analysis (© STRING Consortium 2019). Results are presented as a volcano plot of the significant discriminating functions associated with this principal component. Term related to Lipid oxidation (red), Lipid metabolic processes (blue) and xenobiotics metabolism (yellow) are highlighted as indicated. The highest significant functions belonging to these terms are listed on the right panel. C. Identical genes with a significant Eigen value (> 0.3) and overlapping the indicated GO term were used to build a protein-protein interaction network (© STRING Consortium 2019) with a high level of confidence setting. Genes belonging to a particular organelle network are highlighted in blue (Mitochondria) and red (Peroxysome). Numbers of edges and nodes are indicated as well as PPI enrichment (see methods) D. Identical genes with a significant Eigen value (> 0.3) and overlapping the indicated GO term were used to build a protein-protein interaction network (© STRING Consortium 2019) with a high level of confidence. Genes belonging to a particular network of organelles are highlighted in blue (PPARα specific target genes), green (PPARγ specific target genes) and red (any PPAR target gene). Number of edges and node numbers are indicated as well as PPI enrichment (see methods)

Next, we verified for each treatment dose that the same trends were observed by comparing their RPKM values to those of the control. In this case, we directly used the calculated fold change of genes with an RPKM value > 0.2. Based on an identical approach combining a volcano plot and a protein network reconstruction for each analysis, we demonstrated that genes related to fatty acid oxidation and lipid catabolism were inhibited in treated animals, independently of the doses used (Figure S3A-C and S4A-C, Supplementary Table 6). Of note, significance and enrichment scores of the observed changes increased according to DPhP doses, whereas the density of the protein network encoded by genes altered and related to the lipid catabolism was the highest with the intermediate concentration (edge and node numbers, see methods).

Finally, to exclude a possible artefact arising from poor analytical replicates, we performed a PCA and an OPLS-DA analysis on the 16 analytical samples by comparing each treated group to control animals. Good separation between the replicates was obtained by using the 2^nd^, the 4^th^ and the 5^th^ axis of the PCA (Figure 6A). For each principal component, we retrieved the Eigen values and performed a Gene ontology analysis with these scores (Supplemental Table 7). Using the second axis in which C2 is the most distant from the control, this analysis confirmed the existence of a repression of genes controlling fatty acid catabolism in samples arising from treated animals (Figure 6B). Interestingly, by analysing the two other components, we observed that the highest concentrations of DPhP also increased the expression of genes associated with the synthesis of sterol-derived metabolites, indicating a more complex alteration of lipid metabolism (Figure 6C). Finally, we noticed that the lowest DPhP concentration increased the expression of genes related to the tryptophan and the xenobiotic metabolism (Figure 6D). This was intriguing since tryptophan metabolites have been connected to aryl metabolism and the xenobiotic receptor AHR^28, 29^, whereas our metabolomics approach revealed an accumulation of the same benzene derivatives in a dose-dependent manner. To improve sample clustering, we then tested by OPLS-DA all possible combinations able to discriminate our 4 groups of replicates, using the control replicates as negative controls. Four predictive components were determined (Supplemental Table 8), three of which were plotted as indicated in the Figure 6E and confirmed the reliable separation of the different groups of replicates. The predictive component 3, classifying the samples in a dose/response manner was then used to build a bipartite view of enrichment network associating the 2000 most relevant genes to this component and the KEGG function related to these genes (Supplemental Table 8). The network highlighted two interconnected clusters of genetic programmes repressed in a dose-dependent manner by DPhP, namely the fatty acid metabolism and the xenobiotic response. Inversely, one cluster was related to weakly activated functions belonging to the control of mRNA processes and the acute phase response. The fatty acid cluster associated highly significant programmes related to fatty acid oxidation, peroxysome and PPAR transcriptional response (Figure 6F). Moreover, fatty acid biosynthetic processes such as fatty acid elongation also contributed to this large cluster, these latter functions being inhibited by DPhP treatment. Similarly, the xenobiotic cluster associated several types of xenobiotic responses such as those related to the cytochrome P450, the glucuronidation through the aldarate metabolism, or glutathione metabolism. Lastly, although not present directly in these clusters, we confirmed here that the genetic programme controlling the metabolism of tryptophan was repressed by increasing doses of DPhP and established a clear gene connection (Maob, Hadh, Aldh3a2…) between this response and the two repressed clusters previously mentioned. Of note, results obtained from this last analysis could appear as counterintuitive with regards to the observed activation of tryptophan and xenobiotic responses by the lowest dose of DPhP. Hence, DPhP apparently regulated both responses in a complex inverted U-shape manner with an opposite outcome according to the dose used.

**Figure 6.**
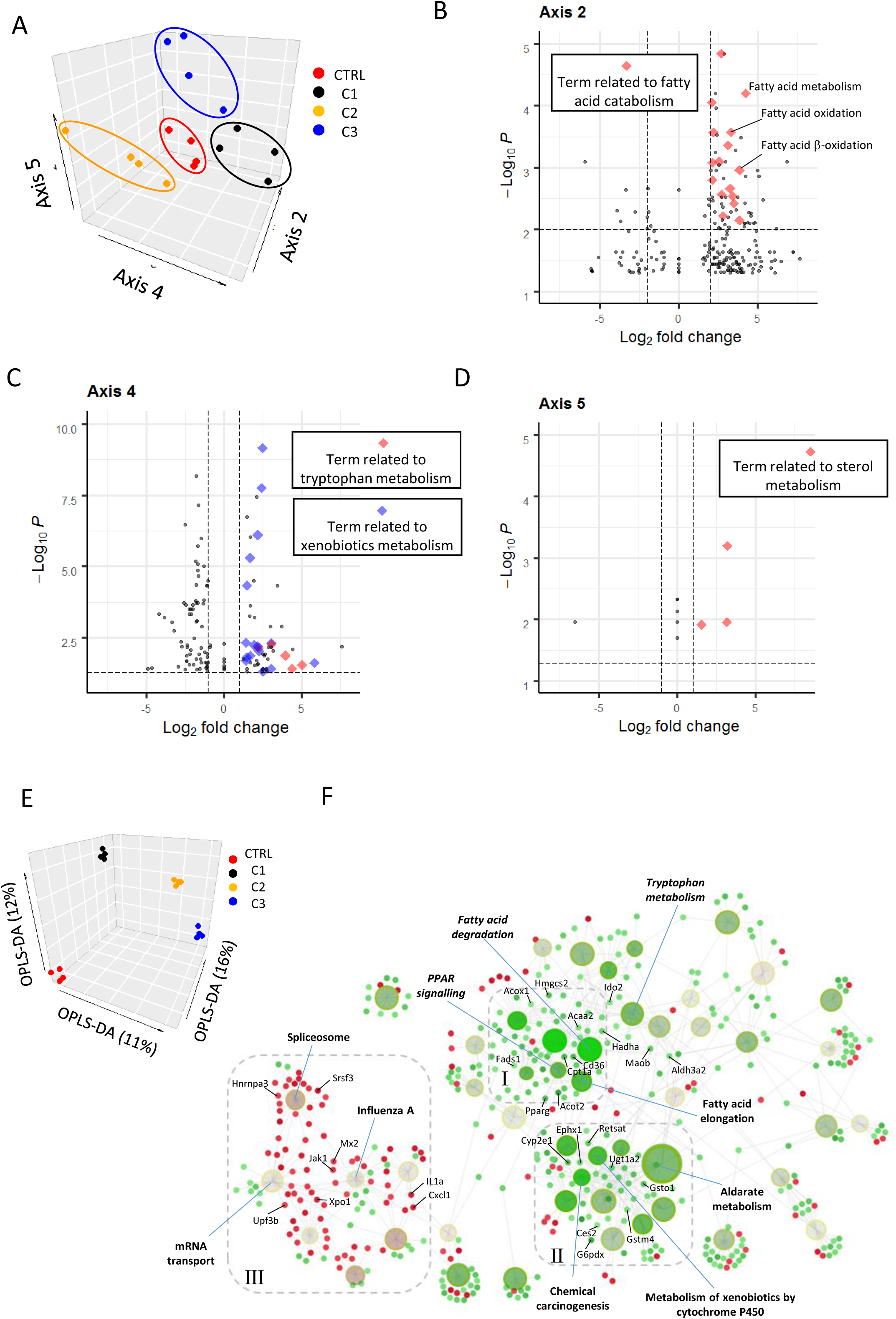
Transcriptomic functions significantly associated with each group of exposed animals A. Principal Component Analysis was performed using pareto algorithm as an observatory method to discriminate individual sample replicates based on the amount of mRNA expression obtained from indicated animals, through reverse transcription and Next-Generation-Sequencing. Individual replicates could be grouped efficiently through the indicated PCA axis. B-D. Gene ontology analysis of the most discriminating genes (Eigen value > 0.3) used to build the indicated axis of the previous PCA performed with STRING. Results are presented as a volcano plot of the significant discriminating function for the indicated axis. Note that axis 2, axis 4 and axis 5 respectively correspond to this order of gene expression, C2<C3<C1<CTL, CTL=C2=C3<C1 and CTL<C1=C2<C3. E. OPLS-DA analysis was performed to discriminate individual sample replicates based on the amount of mRNA expression obtained from indicated animals, through reverse transcription and NGS, and in a supervised manner. A 3D dot plot is presented showing the efficient replicate clustering and the discriminating power attributed to the first 3 predictors. F. Using the predictor (x axis) that classified the samples in a classical dose-response, a bipartite view of enrichment network based on the 2000 most discriminating genes is represented. Small dots and large dots represent individual genes and enriched functions (KEGG based), respectively. Genes or functions are encoded by a green-red scale according to their fold change or their p-value, respectively, as well as the direction (repressed-activated) of the regulation. Dot size of the functions represents the percentage of genes used and matching the full list associated to this function in the KEGG database. 3 clusters of functions are highlighted: I=Lipid metabolism, II=Xenobiotics response, III=mRNA metabolism and acute phase response.

### Chronic DPhP treatment disturbs protein expression of the liver and the overall physiology

Since our multi-omics analysis clearly demonstrated some abnormalities in the processes related to lipid catabolism and PPAR signalling, we performed a series of immunohistochemical analyses revolving around key enzymes of liver physiology, involved in fatty acid catabolic processes and ketone body formation. In addition, we stained the lipid droplets using Perilipin 2 staining, a protein surrounding these vesicles and correlated with their abundance^30^. Furthermore, we analysed enzymes controlling the gluconeogenesis to verify that an overall unspecific dysregulation of liver metabolism did not appear following DPhP treatment (Figure 7A). Expression of Pck1 was thus not repressed by exposure to DPhP, indicating the absence of a global hepatic dysregulation. In some treated animals, Pck1 expression even increased, a mechanism associated with fasting. Conversely, the most striking difference observed was related to the inhibition of protein expression of Hmgcs2, a kernel PPARα target gene, even at the lowest dose of DPhP (Figure 7A). This reduction occurred more specifically in the peri-portal area, where Hmgcs is normally more strongly expressed and active (Figure 7B and S5A). At the highest dose, 100% of animals presented this physiological alteration. Interestingly, this lower expression of HMGCS2 was highly correlated with a reduction of the amount of lipid droplets in animals exposed to 10 mg.L-1 of DPhP (Figure 7A-C and S5B). However, this was not the case for the lowest doses where the amount of Perilipin 2 staining increased upon exposure to DPhP, indicating that lower PPARα and β-oxidative activities may spare some fatty acids from storage depletion in these conditions. PPARα staining was also uncorrelated with Hmgcs2 expression since exposed animals displayed a stronger nuclear and cytosolic expression (Figure 7A), indicating that an active repression of its transcriptional activity was likely taking place after DPhP treatment.

**Figure 7.**
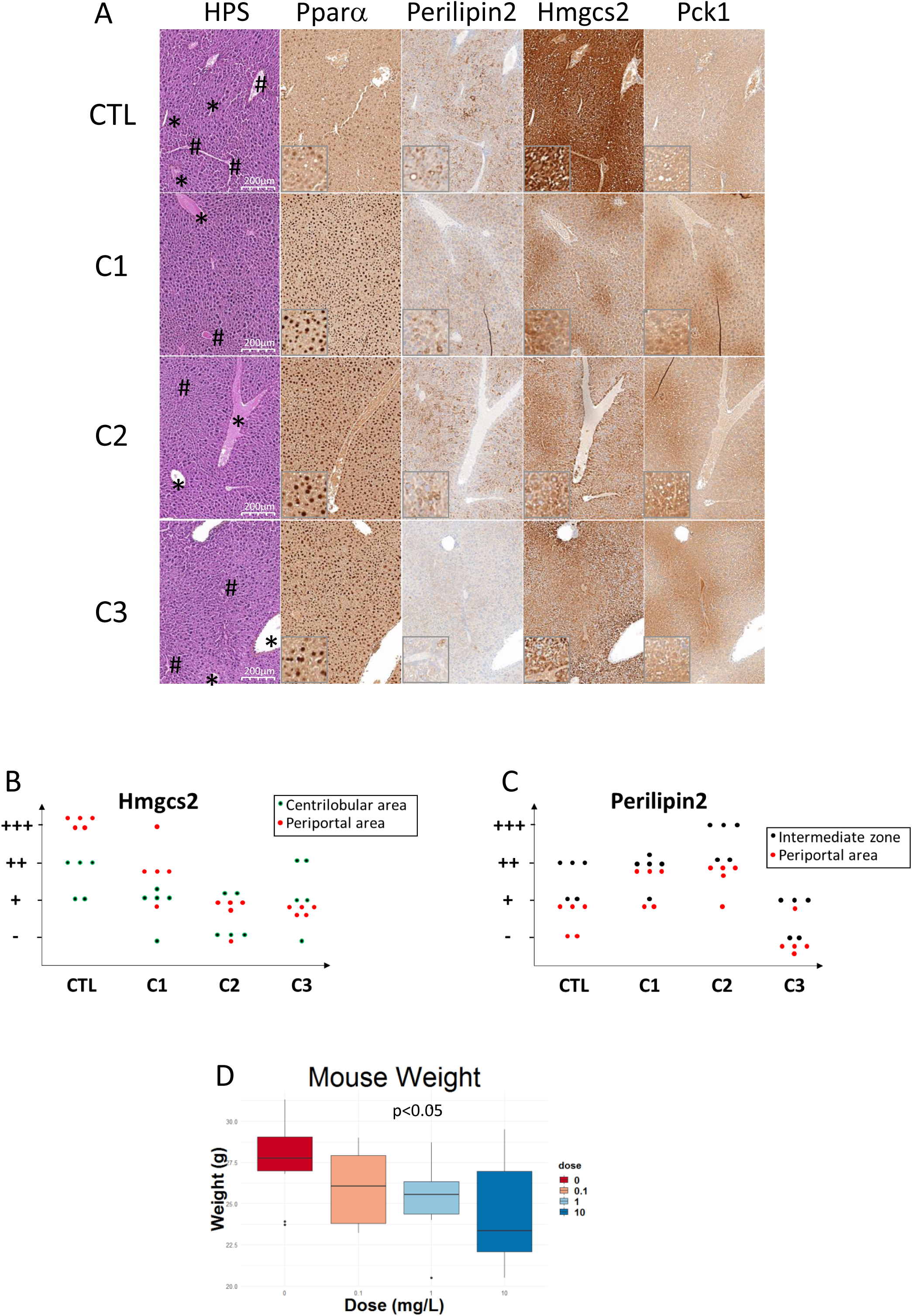
Histological and physiological alterations induced by exposure to DPhP. A. Liver sections at the indicated scale of mice exposed to the indicated concentration of DPhP or a vehicle were immunostained with the indicated antibodies. Star and hash denote the centrolobular and the portal area of the liver, respectively. Inset represents a 10x magnification of the source image. B. Histogram quantifying IHC scores associated with Hmgcs staining of 5 animals in each indicated group (representative of two independent experiments, 2×5 animals). Score associated with the centrilobular and the portal area were dissociated as indicated. (−) = no to very low staining, (+) = low staining, (++) intermediate staining, (+++) = intense staining (see methods). C. Histogram quantifying IHC scores associated with Perilipin 2 staining of 5 animals in each indicated group (representative of two independent experiments, 2×5 animals). Score associated with the intermediate zone and the portal area were dissociated as indicated (no staining was present in the zone contiguous to the centrilobular vein). (−) = no to very low staining, (+) = low staining, (++) intermediate staining, (+++) = intense staining (see methods). D. Box and whisker plot quantifying the body weight of 10 animals representative of two independent experiments (2×5 animals), after 8 weeks of DPhP exposure at the indicated concentrations or with a vehicle. Outliers and significant p-values are indicated.

All of our results highlight the disturbance in lipid metabolism. More specifically, DPhP-treated animals presented signs of fasting, including free fatty acid content and lipid droplet reduction, and reduction of fatty acid biosynthesis and oxidation. We thus used our previous cohort to construct the growth curve of these animals according to their overall weight gain. No significant difference was observed at the lowest doses, but a trend for weight loss was clearly evident in a dose-dependent manner (Figure 7D). Moreover, weight gain was significantly lowered in animals treated with the highest dose of DPhP.

## 3. Discussion

Only few studies have been conducted to directly test the effect of DPhP, the most common APE derivative in human samples. Our results confirm that DPhP levels in biological fluids are unlikely to represent a surrogate of direct APE ingestion, and consequently, are rather a surrogate of their presence in the environment, due to their spontaneous degradation. Therefore, we believe that experimental procedures revolving around DPhP such as those presented in this study are likely more relevant for assessing APE toxicity.

Using this strategy, our results clearly demonstrate that DPhP, even at low doses, disturbs liver metabolism, especially the lipids used in this organ. We clearly observed a reduction of the main free fatty acids (C16.0, C18.0, C18.1) and even a stronger effect on the acylcarnitine pools associated with the degradation of these fatty acids. In line with these results, dodecanedioic acid, a metabolite associated with the impairment of mitochondrial β-oxidation of fatty acids^33^ was dose-dependently accumulated during DPhP exposure. Moreover, our results demonstrate that the genetic programmes associated with the oxidation of fatty acids were strongly repressed, even at the lowest DPhP exposure concentration. Finally, an impact on the liver fat content and body weight of the animals could be observed, especially at the highest dose of DPhP where fatty acids and sterol synthesis was apparently also disturbed in an inverted manner, at least at the transcriptional level. Taken together, these results strongly point towards a significant dysregulation of lipid homeostasis, likely involving PPARα activity and possibly other members of this nuclear receptor family.

PPARα is a key coordinator of fast-fed transition at the hepatic level with paradoxical effects^34^. PPARα is thus the main activator of fatty acid oxidation and ketogenesis during adaptation to long-term fasting^35^, but also in normal conditions to circadian feeding^36^. Therefore, PPARα controls on the one hand *de novo* lipid synthesis and fatty acid uptake during feeding periods, supplying store droplets, and on the other hand it activates triglyceride and cholesteryl-ester lysis and oxidation of released lipids between meals. The simultaneous inhibition of fatty acid oxidation and reduction of the main fatty acid produced by *de novo* synthesis perfectly corresponds to a model of PPARα inhibition upon exposure to DPhP, the lack of fatty acid likely contributing to the overall reduction of acylcarnitines. Our findings that lipid stores in the liver of animals exposed to low concentrations of DPhP are not depleted is also coherent with this model, since an equilibrium may be found between the lack of fatty acid storage and a lower use of these fatty acids. However this may have several consequences, for instance it has been shown that PPARα inhibition is associated with a higher amount of plasma triglyceride and LDL^37, 38^, both being strongly associated with the development of cardiovascular disease, dyslipidemia, metabolic syndrome and even cancer.

Importantly, this equilibrium is likely subtle and not easy to maintain, since we can clearly see a tendency towards weight reduction in exposed animals and a reduction in lipid stores in those ingesting the highest dose of DPhP. Several mechanisms could contribute to these secondary effects, taking place directly in the liver or involving a more complex interaction with other organs. First, despite reduction in fatty acid oxidation, this effect may not be sufficient to compensate for the lack of fatty acid uptake and production at some threshold of PPARα inhibition or after long-term exposure. Second, other members of the PPAR nuclear receptor family may be perturbed, since they are more directly involved in fat digestion, sterol synthesis and storage of fatty acids in the liver and the white adipose tissue^34^. PPARα and γ are thus able to bind the same ligands with a same order of affinity potentially explaining that high doses of DPhP may simultaneously inhibit both types of receptors^39^. PPARγ inhibition in WAT could then be associated with the significant weight loss observed at the highest dose of DPhP. Our observations describing the inhibition of genetic programmes associated with fat digestion and sterol synthesis thus clearly argue in favor of this model.

Interestingly, our study highlights the regulation of xenobiotic and tryptophan metabolism. Indeed, we noticed the alteration of several metabolites, endogenous and exogenous, containing an aromatic ring and/or associated with the tryptophan metabolism. These compounds are implicated in the regulation of AHR, an important xenobiotic receptor^40^. Moreover, even though DPhP does not contain the classical polycyclic structure associated with known ligands of these receptors, it possesses two aromatic rings susceptible to stack with these structures and disturbing their interaction with AHR. Moreover at the genetic level, we also noticed that exposure to DPhP disturbs the xenobiotic response and tryptophan metabolism. We verified the level of AHR, and the expression of canonical target genes of this receptor but did not observe any obvious difference. However, this was expected as mice were not challenged with a classical AHR agonist, although it raises questions as to the possible interactions between DPhP and a classical aromatic polycyclic hydrocarbon. Finally, recent studies suggest that AHR may be involve in fatty acid metabolism^41^, independently of its role as a xenobiotic regulator, potentially explaining part of the biological effects of DPhP.

In conclusion, our results raise many questions on the use, the safety and the presence in the environment of APEs, most of them being susceptible to expose humans to DPhP. We did not fully characterise the molecular mechanisms underlying the apparent alterations in lipid homeostasis and PPAR activities by these compounds, as this was beyond the scope of our study. However, the known function of these factors and their association with the metabolic syndromes should constitute a sufficiently strong risk factor to measure the health hazards associated with their presence in the environment more precisely, by taking into consideration the diet into future epidemiological studies. Moreover, the potential interaction of these compounds with known activators of AHR should be investigated with the aim of determining the possible existence of synergistic effects and to characterise the mechanisms, directly or indirectly, inhibiting PPAR activities.

## 4. Methods

### 4.1. Reagents and chemicals

Diphenyl phosphate, diphenyl phosphate-D10 and triphenyl phosphate-D15 were purchased from Merck (Darmstadt, Germany) with a purity higher than 98%. The confirmation standards of carnitine, acetylcarnitine, palmitoylcarnitine, lauroylcarnitine, decanoylcarnitine, hexanoylcarnitine, stearic acid, oleic acid and linoleic acid were obtained from Merck, all of them with a purity higher than 97%.

Acetonitrile and heptane of LC-MS quality grade and ammonium formate were supplied from by BioSolve (Dieuse, France). Water and formic acid quality grade optima LC-MS were purchased from Fisher Scientific (Illkirch, France).

An emulsion was created containing corn oil for DPhP and TPhP resuspension and animal exposure and 50% of corn oil from Merck (Darmstadt, Germany).

### 4.2. Biological samples from animals and animal care

FVB mice (5 wk old) from Charles River Laboratories were used for all experiments. Animals were housed in the ANICAN (Centre de Recherche en Cancérologie de Lyon) animal facilities accredited by the French Ministry of Agriculture. Food and water were provided *ad libitum* (lights on: 08:00 to 20:00 hours; temperature: 22°C±1°C; humidity: 55%±10%). Experimental groups were designed as follows: a control group and six exposed groups at 0.1 mg/mL (C1), 1 mg/mL (C2) and 10 mg/mL (C3) of DPhP or TPhP for acute exposure. Mice were only treated with DPhP for chronic exposure experiments. At least 5 animals were used in each group for acute exposure for each described series of experiments. For chronic exposure, 2 independent experiments were carried out with 10 animals per group. Metabolomic and transcriptomic analyses were performed on these separate and independent experiments reinforcing the strength of the correlations observed. Between groups, animals were randomized according to their weight at the time of reception. Animal experiments were performed in compliance with French and European regulations on protection of animals used for scientific purposes (EC Directive 2010/63/EU and French Decree 2013–118). They were approved by Ethics Committee and authorized by the French Ministry of Research (APAFIS#3680- 2016010509529577v5).

### 4.3. Sample preparation for metabolomics

A liquid solid extraction was developed for the targeted analysis of diphenyl phosphate and triphenyl phosphate. 20 mg of liver were weighed and 45 µL of internal standard solution at 2 ppm of DPP-d10 and TPP-d15 were added. The mixture was homogenized with a vortex and evaporated. Three zirconium balls, 1 mL of acetonitrile and 0.5 mL of heptane were added. The mixture was homogenized again during 2 min at 3,200 rpm and centrifuged at 10,000 rpm for 7 min. The heptane was discarded, 750 µL of acetonitrile were transferred to a vial and a second extraction with 1mL of acetonitrile was conducted using the same methodology. Finally, all the extracts (total of 1.5 mL of acetonitrile) were pooled, split into two portions of 750 µL, evaporated at 35°C during approximately 90 min and stored at −20°C. The samples were reconstituted with 75 µL of water/acetonitrile 90:10 (*v/v*), prior LC-HRMS analysis reverse phase chromatography, or with 75 µL of acetonitrile/water 95:5 (*v/v*). The extraction was conducted from eight samples of each administered concentration to obtain a suitable, reliable and reproducible statistical results.

### 4.4. UHPLC-ESI-MS/MS analysis

Separation was carried out using an Ultimate 3000 UHPLC system (Thermo Scientific®, MA, USA). The chosen column was a Luna Omega polar C18 (100×2.1 mm, 1.6 µm particle size) (Phenomenex, Torrance, CA, USA), since it has more affinity for polar compounds, to expand the number of detected compounds and detected a larger number of metabolites (normally more polar that parent compounds). A nucleodur HILIC column (100×2.1 mm, 3 µm particle size) (Macherey Nagel, Hoerdt, France) was also used to confirm the identified compounds or detected other compounds that the C18 column did not highlight due to their different polar affinities. The columns were maintained at 30°C during the analysis.

With the C18 column the mobile phase was water/acetonitrile 90/10 (*v/v*) 5 mM ammonium formate and 0.01% formic acid (A) and acetonitrile 5 mM ammonium formate and 0.01% formic acid (B). The gradient elution has a flow of 0.3 mL/min and it started at 100% of A and held during 1 min. The percentage of A then decreased until reaching 0% in 10 min. The gradient was held at this percentage for 4 min prior to finally returning to 100% of A and held for 3 min to condition the column for the next injection. The total running time was 18 min. The injection volume was 5 µL.

For the HILIC column, the mobile phase was water 5 mM ammonium formate and 0.01% formic acid (A) and acetonitrile:water 95/5 (*v/v*) 5 mM ammonium formate and 0.01% formic acid (B). The elution gradient had a flow rate of 0.4 mL/min and started with 95% of B and was held for 2 min. It was then decreased to 70% in 7 min and 50 % in another 2 min. The percentage was held for 4 min to return to the initial percentage in 0.1 min and equilibrated during 10 min. The total running time was 25 min. The injection volume was 5 µL.

The chromatographic system was coupled to a QToF mass spectrometer (Maxis Plus, Bruker Daltonics®, Bremen, Germany) with electrospray ionization interface (ESI) operating in positive and negative mode. The following settings were used: capillary voltage of 3600 V, end plate offset of 500 V, nebulizer pressure of 3 bar (N2), drying gas of 9 L/min (N2), and drying temperature of 200°C. A solution of sodium formate and acetate (10 mM) clusters was used for external calibration at the beginning of each run. The analysis was performed in a full scan over the mass range of 50-1,000 Da with a scan rate of 1 Hz. Moreover, the analysis was carried out in profile mode with the following transfer parameters: funnel 1 RF of 200 Vpp, multipole RF of 50 Vpp, quadrupole energy of 5 eV, collision energy of 7 eV, stepping basic and a pre-pulse storage of 5 ms. The instrument resolution was estimated at 21244 (FWHM) at m/z = 415.211.

The MS/MS experiments were done in data-dependent acquisition mode (AutoMSMS) with a cycle time of 3s and a spectra rate between 2Hz and 16Hz in order to record low and high intensity precursors. They were just conducted on quality control (QC) sample to get MS/MS spectra. Quality controls made by mixing 5 µL of each sample were injected every 13 samples, with a percentage sample/QC approximately of 10%. The samples were analysed randomly to ensure the representativeness of the results.

The software used to acquire data and instrument control were OTOF control 4.1, Hystar 4.1 (Bruker Daltonics®), Data Analysis® 4.4, Metaboscape 4.0 (Bruker Daltonics®) and Mass Frontier^TM^ 7.0 (Thermo Scientific®) were used for data processing.

### 4.5. Annotation workflow

The data obtained from the analysis of the samples were processed using MetaboScape 4.0, including in this analysis all the studied concentrations. The principal parameters used to create the bucket table were: Intensity threshold=5000 counts; minimum peak length=7 spectra; perform MS/MS import, group by collision energy; retention time window (min) =0.4-15; mass window (*m/z*) =60-1000; EIC=0.8; Ions=H^+^, Na^+^, K^+^, NH4^+^, −H2O+H^+^. The bucket table (1720 couples or features) contained information regarding retention time, ion *m/z* ratio, neutral mass, detected ions, MS/MS spectrum and relative peak intensity of features in each sample.

The bucket tables were analysed by unsupervised principal components analysis (PCA) conducted simultaneously on all samples (C1, C2, C3 and control), using the pareto algorithm as an observatory method to discriminate groups of samples. Supervised partial least squares (PLS) (using pareto algorithm) and T-Test (using group mean algorithm), individually comparing each contamination concentration and control were conducted. A *p-*value < 0.05 was considered to be significantly differentially expressed.

For compound annotations, the formulas of the couples with a *p*-value < 0.05 were selected based on a mass deviation under 5 mDa and 2 ppm. mSigma (comparison between theoretical and experimental isotopic pattern) less than 20, was also taken into consideration when searching for compounds. The discovered formulas were from different databases (analyte DB, ChEBI, ChemSpider, PubChem, HMDB) and the compounds with a logical biological link with the matrix (compounds naturally present in liver or biological tissues) or previously detected in biological samples according to the literature were annotated for posterior evaluation. This process was done using the tool compound crawler, included in MetaboScape 4.0. Known intermediate metabolites were also investigated manually to complete the network according to a mass deviation less than 50 mDa and 20 ppm. Absence of significant p-value between conditions was not considered in this setting.

### 4.6. Putative identification of metabolites

Some of the annotated masses had MS/MS spectra. The theoretical formulas for these masses were *in silico* fragmented using MetFrag (MetaboScape) and Mass Frontier. The compounds with a correlation of fragments higher than a 90% were annotated as putatively identified. The HILIC column was then used to compare the retention time obtained with both columns and to search new discriminant compounds through supervised multivariate statistical analysis (PLS and T-Test). Finally, in order to confirm these results, the analytical standards of some of putative compounds with a logical retention time in both columns were purchased. The spiked samples were those with the detected compounds had MS/MS spectra or with the highest concentration in the detected compounds. Besides, both were analysed, spiked and non-spiked samples to see the intensity differences between them in case retention times were the same. Comparison of MSMS spectra of spiked and non-spiked samples was performed to confirm the compounds.

### 4.7 Chemrich and MetaMapp annotation

KEGG ID, Pubchem ID, SMILE and InchiKeys of annotated metabolites were retrieved from public databases as indicated in the former publication of these statistical tools. P-value and fold enrichment were obtained from the signal intensities associated with the multiple injections of each group of samples. Chemrich analyses were then obtained directly from the Chemrich interface website, whereas MetaMapp results were loaded in the cytoscape software to represent the network with an organic layout. Cluster name were retrieved in an unbiased way from Chemrich results tables. All tables associated with these analyses are available in the supplemental information section.

### 4.8. Sample preparation for transcriptomic analyses and Next-Generation Sequencing

#### RNA extraction for tissue

Total RNA was extracted and purified using the RNeasy Mini Kit (Qiagen # 74106) from mouse liver. Furthermore, the RNeasy procedure enriches RNA species > 200 nt and excludes 5S rRNA, tRNAs, or other low molecular weight RNAs. RNA was isolated on the silica membrane in trusted RNeasy spin columns, with binding capacities of 100 μg of RNA, according to the supplier’s recommendations (Qiagen).

#### RNAseq library sequencing and analysis

For the preparation of the NGS RNA library, RNA concentration was measured using the GE NanoView Spectrophotometer (Biochrom US, Holliston, MA, US). The quality of RNA samples was analyzed using the RNA 6000 Pico Kit running on the 2100 BioAnalyzer (Agilent Santa Clara, California, US). Total RNA was diluted in a final volume of 50 μL for a total input of 1 μg. Only the RNA pools with a RIN score higher than 7 were used in the NGS library preparation prior to sequencing.

Firstly, mRNAs were isolated using the NEBNext Poly(A) mRNA Magnetic Isolation Module from 1 μg of total RNA, in triplicate for each condition. The isolation procedure is based on the selection of mRNA using oligo dT beads directed against polyA tails of intact mRNA. Secondly, the NGS libraries were created from mRNA isolated using the NEBNext Ultra II Directional RNA Library Prep Kit for Illumina (NewEngland BioLabs, Ipswich, Massachusetts). Final libraries were sequenced on a Illumina NextSeq 500 on a high output flowcell with 2×75 bp paired-end read lengths.

The sequencing reads were obtained after demultiplexing the raw sequencing data using bcl2fastq v2.19.1.403 (Version v2.15.0 for NextSeq™ 500 and HiSeq® X Systems, Illumina), after having validated the quality controls of each sample using the FastQC v0.11.5 software (https://www.bioinformatics.babraham.ac.uk/projects/fastqc/). The alignment files were generated with STAR v2.5.2b (University of Birmingham) in the 2-pass mode. We used the GRCm38, version M16 (Ensembl 91) as reference. This mode is known to improve the detection of more reads mapping novel splice junctions.

### 4.9. Gene ontology and statistical analyses of NGS

Genes were filtered through their obtained RPKM. Only genes sufficiently expressed in control conditions were conserved for further analyses (RPKM > 0.2). PCA and Orthogonal Projections to Latent Structures Discriminant Analysis were performed on the R platform, using ROPLS package and 3D plots for visualization of the results. Gene ontology analyses were performed by using the Eigen values associated with the most significant axis used to construct PCA and OPLS-DA. With these methods, all conditions, the three different DPhP concentrations and the vehicle, were thus taken into consideration and simultaneously compared. Threshold for retained Eigen values used to select genes used in these analyses are indicated directly in the figures. Functional Enrichment Analysis (© STRING Consortium 2019) was then performed by sorting the selected genes according to their Eigen values. A volcano plot (R package EnhanceVolcano Plot) was eventually used to represent the discriminating functions for the studied axis through the enrichment score and the false discovery rate generated by the STRING algorithm. Only significant functions were retrieved.

Alternatively, paired analyses were performed using each concentration of DPhP against the vehicle. In this case, retained genes were selected through the fold change between both conditions. Gene ontology analysis was then performed identically with Functional Enrichment Analysis (© STRING Consortium 2019).

Finally, in a complementary approach, density of the protein networks associated with discriminating functions and encoded by the genes selected with these different approaches were measured through protein-protein association networks (© STRING Consortium 2019). High number of nodes and edges indicate that large functional complexes are disturbed through multiple genetic regulations induced by DPhP exposure, reinforcing the probability of that function being disturbed by this condition. We used high-confidence settings to retain the experimentally validated interactions. A protein-protein interaction enrichment p-value was calculated against an identical number of random proteins.

### 4.10. Bipartite view of enrichment network

NetworkAnalysis server (Mc Gill University) was used through the List Enrichment function. Functions belonging to the KEGG database were used and only those with a p.value < 0.03 were conserved. Metabolic pathways and carbon metabolism functions were removed due to their very high coverage of the genome. The bipartite view was then selected and an auto-layout was applied. Clustering between functions was then enhanced through the Force Atlas tool. Clusters were then manually highlighted and default encoding of the functions through their p-value was converted into a continuous green-red scale.

### 4.11. Immunohistochemistry analyses

For histological examination, tissue samples were fixed in 10% buffered formalin and embedded in paraffin. 4-µm-thick tissue sections of formalin-fixed, paraffin-embedded tissue were prepared according to conventional procedures. Sections were then stained with haematoxylin and eosin and examined under a light microscope.

Immunohistochemistry was performed on an automated immunostainer (Ventana Discovery XT, Roche, Meylan, France) using Omnimap DAB Kit according to the manufacturer’s instructions. Sections were incubated with the following antibodies: Anti-Perilipin2 (AtlasAntibodies-HPA016607), Anti-Hmgcs2 (Santa Cruz-sc-376092), Anti-PPARalpha (Abnova-MAB12349), Anti-Pck1 (AtlasAntibodies-HPA006507), Anti-FBP1 (Abcam- ab109020) (all diluted at 1:100). An anti-rabbit/mouse - HRP was applied on sections. Staining was visualized with DAB solution with 3,3-diaminobenzidine as a chromogenic substrate.

Sections were counterstained with Gill’s haematoxylin and finally scanned with panoramic scan II (3D Histech, Budapest, Hungary) at 20X. Scoring was performed by three independent investigators using a staining scale ranging from 0 to 10. The means were then calculated and encoded as follows. 0-2=no staining to very low staining (−), 2-4=low staining (+), 4-7=intermediate staining (++), 7-10=high staining (+++).

## ACKNOWLEDGMENTS

The authors would like to thank N. Gadot for IHC analyses, I. Puisieux for its support to design animal experiments and B. Manship for critical reading of the manuscript. This study war strongly financed by the Plan Cancer et Environnement 2015-2019 (project PLASTOC-Plastic additives: study Of the potential risk for Cancer development through the evaluation of pharmacodynamics, accumulation and mechanisms in endocrine tissues). Our team is also certified by the Ligue nationale contre le cancer. Additionally, this work was supported by the Ligue contre le cancer (Comité du Rhône), the Association pour la Recherche contre le Cancer (project n° SFI20121205395,PJA20141201758, PJA 20171206303), and the Bonus Qualité Recherche (BQR) Accueil of Lyon University.

## AUTHOR CONTRIBUTIONS

SR, MA and JD performed experiments on cell lines and animal models. SR coordinated animal experimentation and IHC analysis. JG, CB and LPG performed NGS and their bioinformatics analysis. AMV performed statistical retreatment of the data. JMS, AF, AB and EV designed and supervised metabolomics analyses and raw data retreatment. JMS and AMV performed then their statistical interpretation. LS and BG performed supervised quantification of APEs. EV supervised these analyses. BF, SI and TG helped to interpret the data. LPG and AMV conceived the project, designed experiments, interpreted data, and wrote the manuscript. All authors read and approved the final version of the manuscript.

## References

1. Ballesteros-Gomez, A. et al. In vitro metabolism of 2-ethylhexyldiphenyl phosphate (EHDPHP) by human liver microsomes. Toxicol Lett 232, 203–12 (2015).

2. Ballesteros-Gomez, A., Van den Eede, N. & Covaci, A. In vitro human metabolism of the flame retardant resorcinol bis(diphenylphosphate) (RDP). Environ Sci Technol 49, 3897–904 (2015).

3. Heitkamp, M.A., Freeman, J.P., McMillan, D.C. & Cerniglia, C.E. Fungal metabolism of tert-butylphenyl diphenyl phosphate. Appl Environ Microbiol 50, 265–73 (1985).

4. Bjornsdotter, M.K. et al. Presence of diphenyl phosphate and aryl-phosphate flame retardants in indoor dust from different microenvironments in Spain and the Netherlands and estimation of human exposure. Environ Int 112, 59–67 (2018).

5. Fu, L. et al. Organophosphate Triesters and Diester Degradation Products in Municipal Sludge from Wastewater Treatment Plants in China: Spatial Patterns and Ecological Implications. Environ Sci Technol 51, 13614–13623 (2017).

6. Li, J. et al. A review on organophosphate Ester (OPE) flame retardants and plasticizers in foodstuffs: Levels, distribution, human dietary exposure, and future directions. Environ Int 127, 35–51 (2019).

7. Wang, Y. & Kannan, K. Concentrations and Dietary Exposure to Organophosphate Esters in Foodstuffs from Albany, New York, United States. J Agric Food Chem 66, 13525–13532 (2018).

8. Wang, Y., Kannan, P., Halden, R.U. & Kannan, K. A nationwide survey of 31 organophosphate esters in sewage sludge from the United States. Sci Total Environ 655, 446–453 (2019).

9. Su, G., Letcher, R.J. & Yu, H. Organophosphate Flame Retardants and Plasticizers in Aqueous Solution: pH-Dependent Hydrolysis, Kinetics, and Pathways. Environ Sci Technol 50, 8103–11 (2016).

10. Mitchell, C.A. et al. Diphenyl Phosphate-Induced Toxicity During Embryonic Development. Environ Sci Technol 53, 3908–3916 (2019).

11. Jamarani, R., Erythropel, H.C., Nicell, J.A., Leask, R.L. & Maric, M. How Green is Your Plasticizer? Polymers (Basel) 10(2018).

12. Meeker, J.D., Cooper, E.M., Stapleton, H.M. & Hauser, R. Urinary metabolites of organophosphate flame retardants: temporal variability and correlations with house dust concentrations. Environ Health Perspect 121, 580–5 (2013).

13. Meeker, J.D. & Stapleton, H.M. House dust concentrations of organophosphate flame retardants in relation to hormone levels and semen quality parameters. Environ Health Perspect 118, 318–23 (2010).

14. Sasaki, K., Suzuki, T., Takeda, M. & Uchiyama, M. Metabolism of phosphoric acid triesters by rat liver homogenate. Bull Environ Contam Toxicol 33, 281–8 (1984).

15. Su, G., Crump, D., Letcher, R.J. & Kennedy, S.W. Rapid in vitro metabolism of the flame retardant triphenyl phosphate and effects on cytotoxicity and mRNA expression in chicken embryonic hepatocytes. Environ Sci Technol 48, 13511–9 (2014).

16. Zhao, F., Kang, Q., Zhang, X., Liu, J. & Hu, J. Urinary biomarkers for assessment of human exposure to monomeric aryl phosphate flame retardants. Environ Int 124, 259–264 (2019).

17. Van den Eede, N., Ballesteros-Gomez, A., Neels, H. & Covaci, A. Does Biotransformation of Aryl Phosphate Flame Retardants in Blood Cast a New Perspective on Their Debated Biomarkers? Environ Sci Technol 50, 12439–12445 (2016).

18. Hou, R. et al. Enhanced degradation of triphenyl phosphate (TPHP) in bioelectrochemical systems: Kinetics, pathway and degradation mechanisms. Environ Pollut 254, 113040 (2019).

19. Abdallah, M.A., Nguyen, K.H., Moehring, T. & Harrad, S. First insight into human extrahepatic metabolism of flame retardants: Biotransformation of EH-TBB and Firemaster-550 components by human skin subcellular fractions. Chemosphere 227, 1–8 (2019).

20. Mendelsohn, E. et al. Nail polish as a source of exposure to triphenyl phosphate. Environ Int 86, 45–51 (2016).

21. Chen, G., Jin, Y., Wu, Y., Liu, L. & Fu, Z. Exposure of male mice to two kinds of organophosphate flame retardants (OPFRs) induced oxidative stress and endocrine disruption. Environ Toxicol Pharmacol 40, 310–8 (2015).

22. Wang, D. et al. Neonatal triphenyl phosphate and its metabolite diphenyl phosphate exposure induce sex- and dose-dependent metabolic disruptions in adult mice. Environ Pollut 237, 10–17 (2018).

23. Belcher, S.M., Cookman, C.J., Patisaul, H.B. & Stapleton, H.M. In vitro assessment of human nuclear hormone receptor activity and cytotoxicity of the flame retardant mixture FM 550 and its triarylphosphate and brominated components. Toxicol Lett 228, 93–102 (2014).

24. Tung, E.W.Y., Ahmed, S., Peshdary, V. & Atlas, E. Firemaster(R) 550 and its components isopropylated triphenyl phosphate and triphenyl phosphate enhance adipogenesis and transcriptional activity of peroxisome proliferator activated receptor (Ppargamma) on the adipocyte protein 2 (aP2) promoter. PLoS One 12, e0175855 (2017).

25. Kojima, H. et al. In vitro endocrine disruption potential of organophosphate flame retardants via human nuclear receptors. Toxicology 314, 76–83 (2013).

26. Barupal, D.K. & Fiehn, O. Chemical Similarity Enrichment Analysis (ChemRICH) as alternative to biochemical pathway mapping for metabolomic datasets. Sci Rep 7, 14567 (2017).

27. Barupal, D.K. et al. MetaMapp: mapping and visualizing metabolomic data by integrating information from biochemical pathways and chemical and mass spectral similarity. BMC Bioinformatics 13, 99 (2012).

28. Venkateswaran, N. et al. MYC promotes tryptophan uptake and metabolism by the kynurenine pathway in colon cancer. Genes Dev 33, 1236–1251 (2019).

29. Yamamoto, T. et al. Kynurenine signaling through the aryl hydrocarbon receptor maintains the undifferentiated state of human embryonic stem cells. Sci Signal 12(2019).

30. Straub, B.K. et al. Adipophilin/perilipin-2 as a lipid droplet-specific marker for metabolically active cells and diseases associated with metabolic dysregulation. Histopathology 62, 617–31 (2013).

31. Su, G., Letcher, R.J., Yu, H., Gooden, D.M. & Stapleton, H.M. Determination of glucuronide conjugates of hydroxyl triphenyl phosphate (OH-TPHP) metabolites in human urine and its use as a biomarker of TPHP exposure. Chemosphere 149, 314–9 (2016).

32. Roman, P., Cardona, D., Sempere, L. & Carvajal, F. Microbiota and organophosphates. Neurotoxicology 75, 200–208 (2019).

33. Korman, S.H., Waterham, H.R., Gutman, A., Jakobs, C. & Wanders, R.J. Novel metabolic and molecular findings in hepatic carnitine palmitoyltransferase I deficiency. Mol Genet Metab 86, 337–43 (2005).

34. Dubois, V., Eeckhoute, J., Lefebvre, P. & Staels, B. Distinct but complementary contributions of PPAR isotypes to energy homeostasis. J Clin Invest 127, 1202–1214 (2017).

35. Janssen, A.W. et al. The impact of PPARalpha activation on whole genome gene expression in human precision cut liver slices. BMC Genomics 16, 760 (2015).

36. Gachon, F. et al. Proline- and acidic amino acid-rich basic leucine zipper proteins modulate peroxisome proliferator-activated receptor alpha (PPARalpha) activity. Proc Natl Acad Sci U S A 108, 4794–9 (2011).

37. Schoonjans, K. et al. PPARalpha and PPARgamma activators direct a distinct tissue-specific transcriptional response via a PPRE in the lipoprotein lipase gene. EMBO J 15, 5336–48 (1996).

38. Watts, G.F. et al. Differential regulation of lipoprotein kinetics by atorvastatin and fenofibrate in subjects with the metabolic syndrome. Diabetes 52, 803–11 (2003).

39. Lim, H. & Dey, S.K. A novel pathway of prostacyclin signaling-hanging out with nuclear receptors. Endocrinology 143, 3207–10 (2002).

40. Bock, K.W. Aryl hydrocarbon receptor (AHR): From selected human target genes and crosstalk with transcription factors to multiple AHR functions. Biochem Pharmacol 168, 65–70 (2019).

41. Bock, K.W. Aryl hydrocarbon receptor (AHR) functions in NAD(+) metabolism, myelopoiesis and obesity. Biochem Pharmacol 163, 128–132 (2019).

