## Supplemental Data for "*In vivo* characterisation of the toxicological properties of DPhP, one of the main degradation products of aryl phosphate esters"

### Supplemental Material

***In vivo* characterisation of toxicological properties related to DPhP, a principal degradative product of aryl phosphate esters.**

Samia Ruby[1]*, Jesús Marín-Sáez[2]*, Aurélie Fildier[3]*, Audrey Buleté[3], Myriam Abdallah[1], Jessica Garcia[4], Julie Deverchère[1], Loïc Spinner[3], Barbara Giroud[3], Sébastien Ibanez[5], Thierry Granjon[5], Claire Bardel[1], Béatrice Fervers[6], Alain Puisieux[1], Emmanuelle Vulliet[3], Léa Payen[1,4]†, Arnaud Vigneron[1]†

#### Table of Contents

**Figure S1.** Annotation strategy and MS/MS validation of critical compounds.

A. Signal intensity of most significant compounds were retrieved. A box and whisker plot could then be constructed to validate their higher or lower presence in exposed animals. An example is shown for butyryl-Lcarnitine.

B. Compound annotation with some proposals of formulas. The first one corresponding to butyryl-L-carnitine.

C. An exemple of fragmentation prediction is shown for this same butyryl-L-carnitine

D. MSMS experimental spectra of butyryl-L-carnitine

**Figure S2.** Metabolic network analysis through structural identity and pathway mapping

A-B. ChemRICH set enrichment statistics plot for 175 metabolites extracted from exposed animals with the indicated concentration of DPhP versus control. Each node reflects a significantly altered cluster of metabolites. Enrichment p-values are given by the Kolmogorov–Smirnov-test. Node sizes represent the total number of metabolites in each set of clusters. The node colour scale shows the proportion of increased (red) or decreased (blue) compounds in exposed mice compared to control mice. Purple-colour nodes have both increased and decreased metabolites.

C-D. MetaMapp visualization of the same metabolomics data highlighting the differential metabolic regulation and the organization of metabolic clusters based on KEGG reactant pair information and Tanimoto chemical similarity matrix.  Increased metabolite levels in exposed mice are labelled as red nodes, decreased levels are marked in blue. Intensity difference are also encoded in node size. Cluster label is indicated. Metabolites previously clustered together based on their structural similarity can now be separated according to different pathway mapping (dinucleotides are now present in the tryptophan metabolism cluster, whereas purine are in the nuclei base metabolism).

**Figure S3**. Gene ontology analysis of transcripts significantly dysregulated by exposure to DPhP

A-C. Gene ontology analysis using the Log2 Fold Change of mRNA expression derived from animals exposed to the indicated concentration of DPhP vs control animals and performed through Functional Enrichment Analysis (© STRING Consortium 2019). Results are presented as a volcano plot of the significant discriminating functions. Term related to Lipid oxidation, Lipid metabolic processes and xenobiotics metabolism are highlighted as indicated.

**Figure S4.** Protein interaction network based on significantly dysregulated genes by exposure to DPhP.

A-C. Genes significantly dysregulated by an exposure to the indicated concentrations of DPhP and overlapping the indicated GO term were used to build a protein-protein interaction network (© STRING Consortium 2019) with a high level of confidence. Genes belonging to a particular network are highlighted in blue (Mitochondria), red (Peroxysome) or green (glycerophospholipid). Numbers of edges and nodes are indicated, as well as PPI enrichment (see methods).

**Figure S5.** Histological alterations induced by exposure to DPhP

A. Liver sections of mice exposed to the indicated concentrations of DPhP or a vehicle were immunostained with an antibody directed against Hmgcs2. Star, cross and hash denote the centrilobular, the intermediate and the portal area of the liver, respectively.

B. Liver sections of mice exposed to the indicated concentrations of DPhP or a vehicle were immunostained with an antibody directed against Perilipin 2. Star, cross and hash denote the centrilobular, the intermediate and the portal area or the liver, respectively.

- Supplementary Figures

A

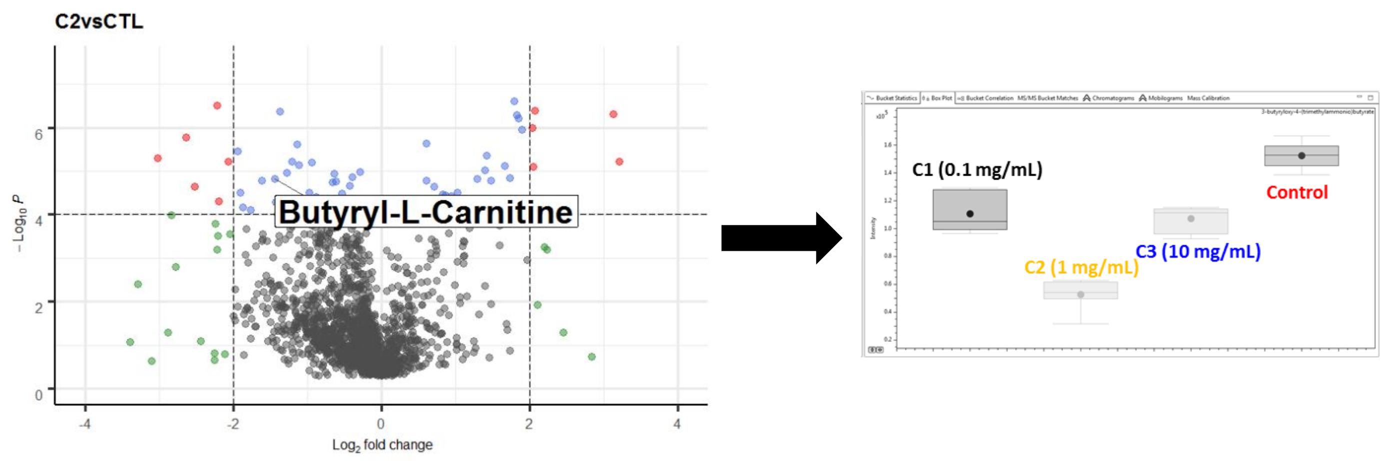


B

| # | Neutral Formula | Ions | Ion Formula | M calc. | Ion | m/z m... | m/z calc. | rdb ... | eConf | Δ m/z [mDa] | Δ m/z [ppm] | mSigma | # Frag | IntCov. [%] |
|---|---|---|---|---|---|---|---|---|---|---|---|---|---|---|
| 1 | $C_{11}H_{21}NO_4$ | ▫ | $C_{11}H_{22}NO_4^+$ | 231.1471 | [M+H]+ | 232.1545 | 32.15433461 | 2.0 | Even | -0.1565 | -0.67 | 2.96 | 5 | 96.9 |
| 2 | $C_{12}H_{17}N_5$ | ▫ | $C_{12}H_{18}N_5^+$ | 231.1484 | [M+H]+ | 232.1545 | 32.15567202 | 7.0 | Even | 1.1810 | 5.09 | 10.76 | 3 | 13.1 |
| 3 | $C_9H_{21}N_5S$ | ▫ | $C_9H_{22}N_5S^+$ | 231.1518 | [M+H]+ | 232.1545 | 32.15904333 | 4.0 | Even | 4.5523 | 19.61 | 30.70 | 3 | 13.1 |
| 4 | $C_7H_{17}N_7O_2$ | ▫ | $C_7H_{18}N_7O_2^+$ | 231.1444 | [M+H]+ | 232.1545 | 32.15164927 | 3.0 | Even | -2.8418 | -12.24 | 32.22 | 5 | 96.9 |

C

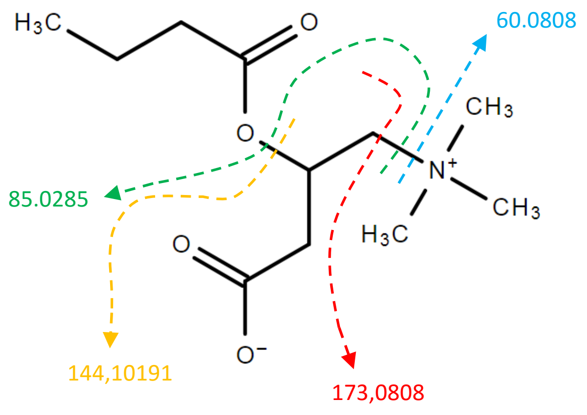


O-butanoylcarnitine

D

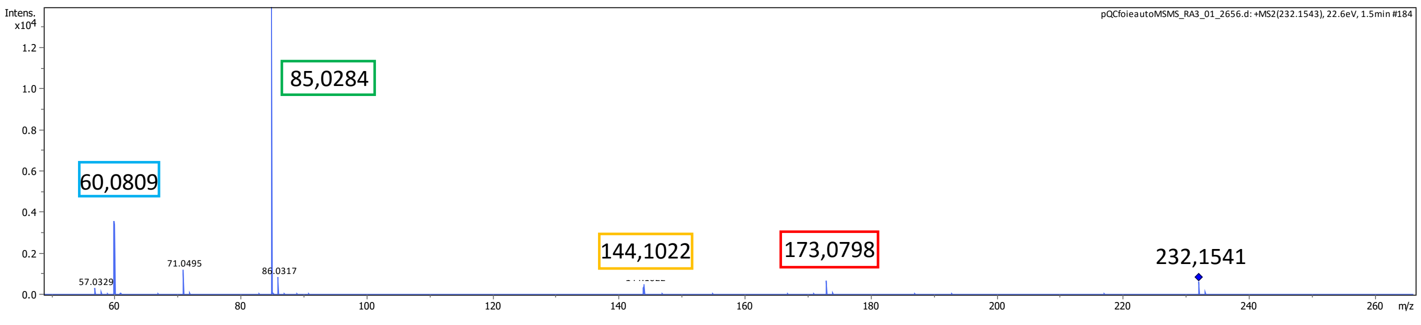

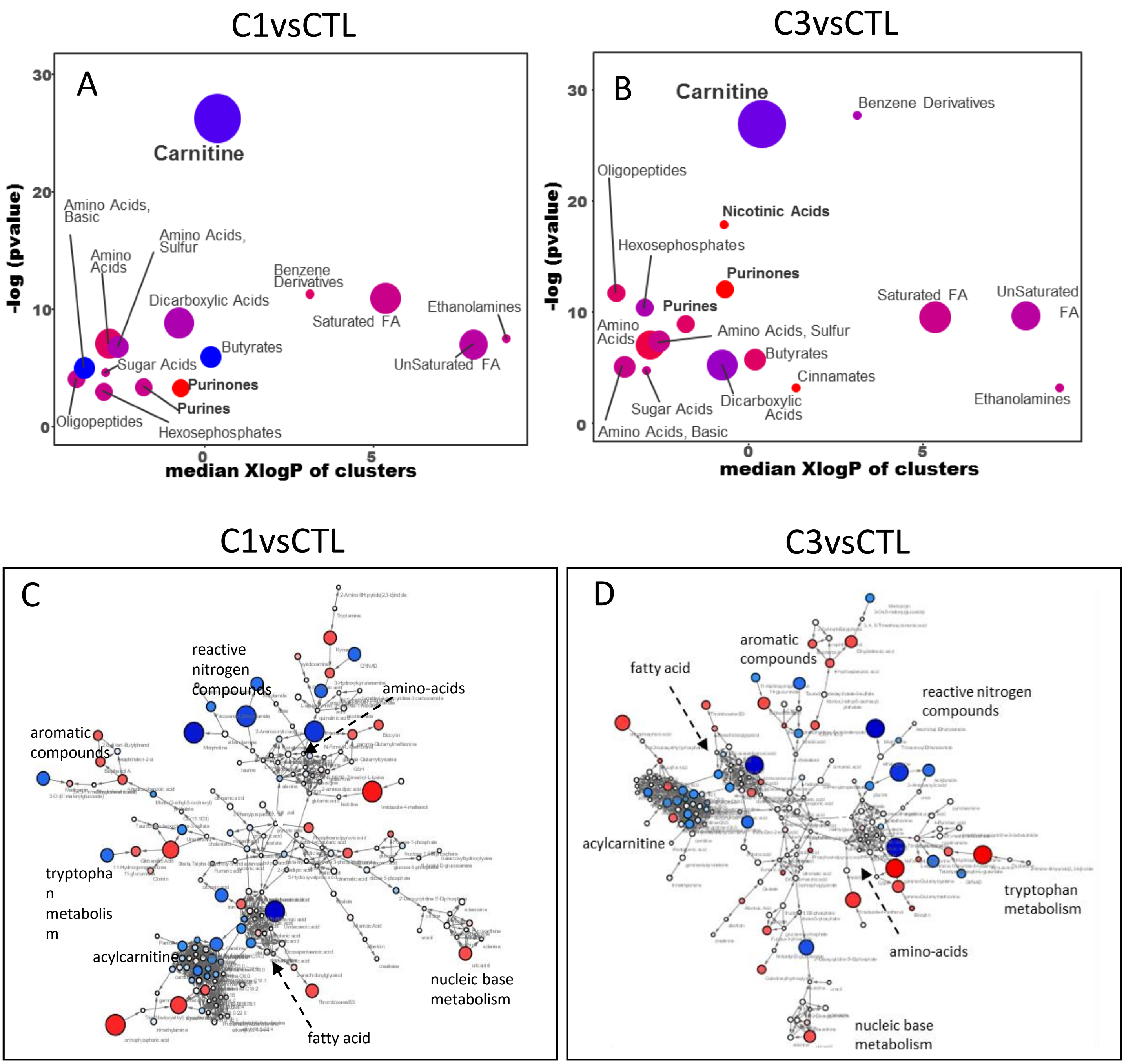
C1vsCTL
C3vsCTL
A
Carnitine
Amino Acids, Basic
Amino Acids, Sulfur
Amino Acids
Benzene Derivatives
Dicarboxylic Acids
Saturated FA
Ethanolamines
Butyrates
Sugar Acids
UnSaturated FA
Purinones
Purines
Oligopeptides
Hexosephosphates
-log (pvalue)
median XlogP of clusters
B
Carnitine
Benzene Derivatives
Oligopeptides
Nicotinic Acids
Hexosephosphates
Purinones
Purines
Saturated FA
UnSaturated FA
Amino Acids
Amino Acids, Sulfur
Butyrates
Cinnamates
Sugar Acids
Dicarboxylic Acids
Ethanolamines
Amino Acids, Basic
C1vsCTL
C3vsCTL
C
reactive nitrogen compounds
amino-acids
aromatic compounds
tryptophan metabolism
acylcarnitine
nucleic base metabolism
fatty acid
D
aromatic compounds
fatty acid
reactive nitrogen compounds
acylcarnitine
tryptophan metabolism
amino-acids
nucleic base metabolism

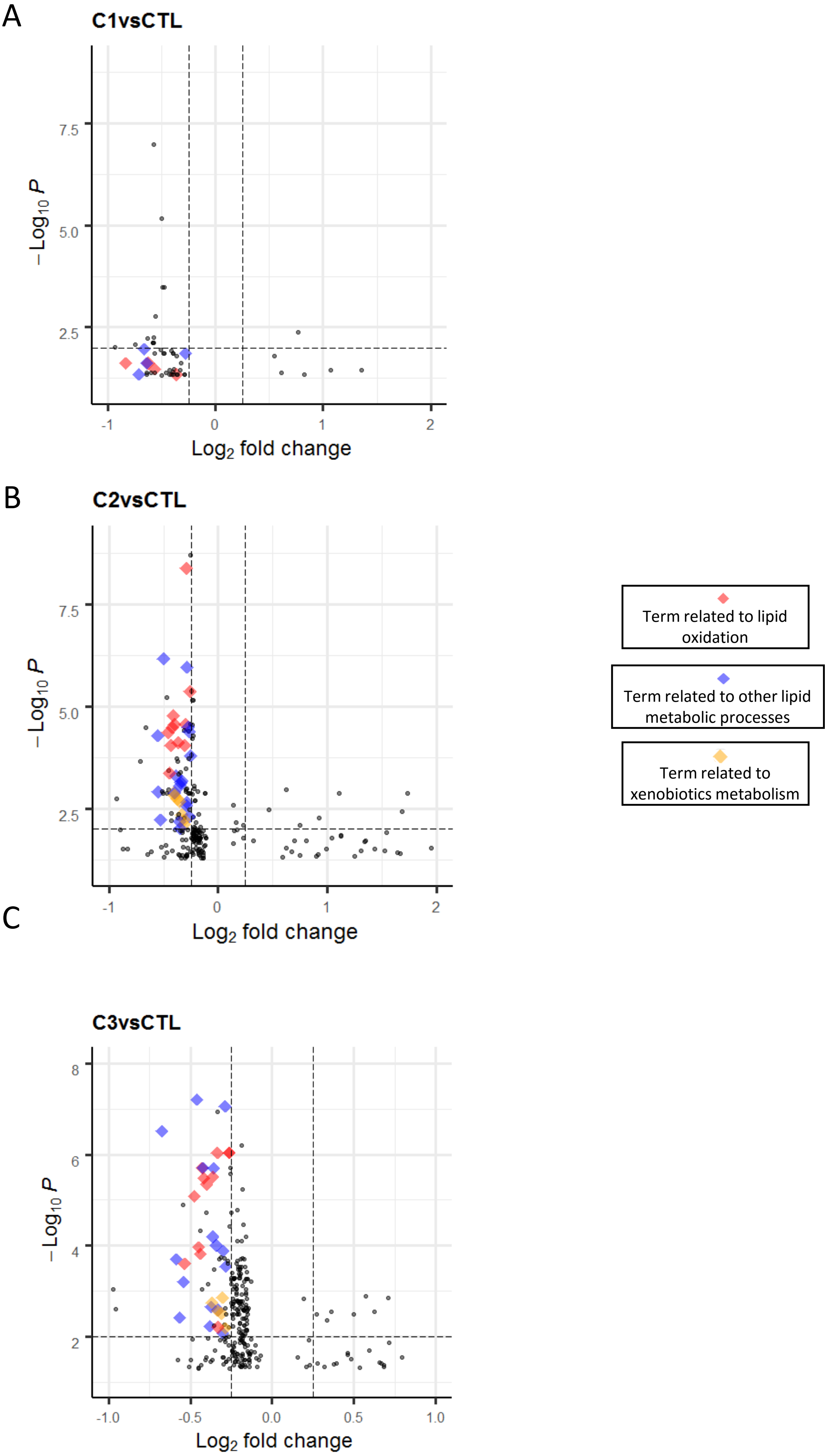
A
C1vsCTL
7.5
5.0
2.5
−Log$_{10}$ P
-1
0
1
2
Log$_2$ fold change
B
C2vsCTL
7.5
5.0
2.5
−Log$_{10}$ P
-1
0
1
2
Log$_2$ fold change
C
C3vsCTL
8
6
4
2
−Log$_{10}$ P
-1.0
-0.5
0.0
0.5
1.0
Log$_2$ fold change
Term related to lipid oxidation
Term related to other lipid metabolic processes
Term related to xenobiotics metabolism

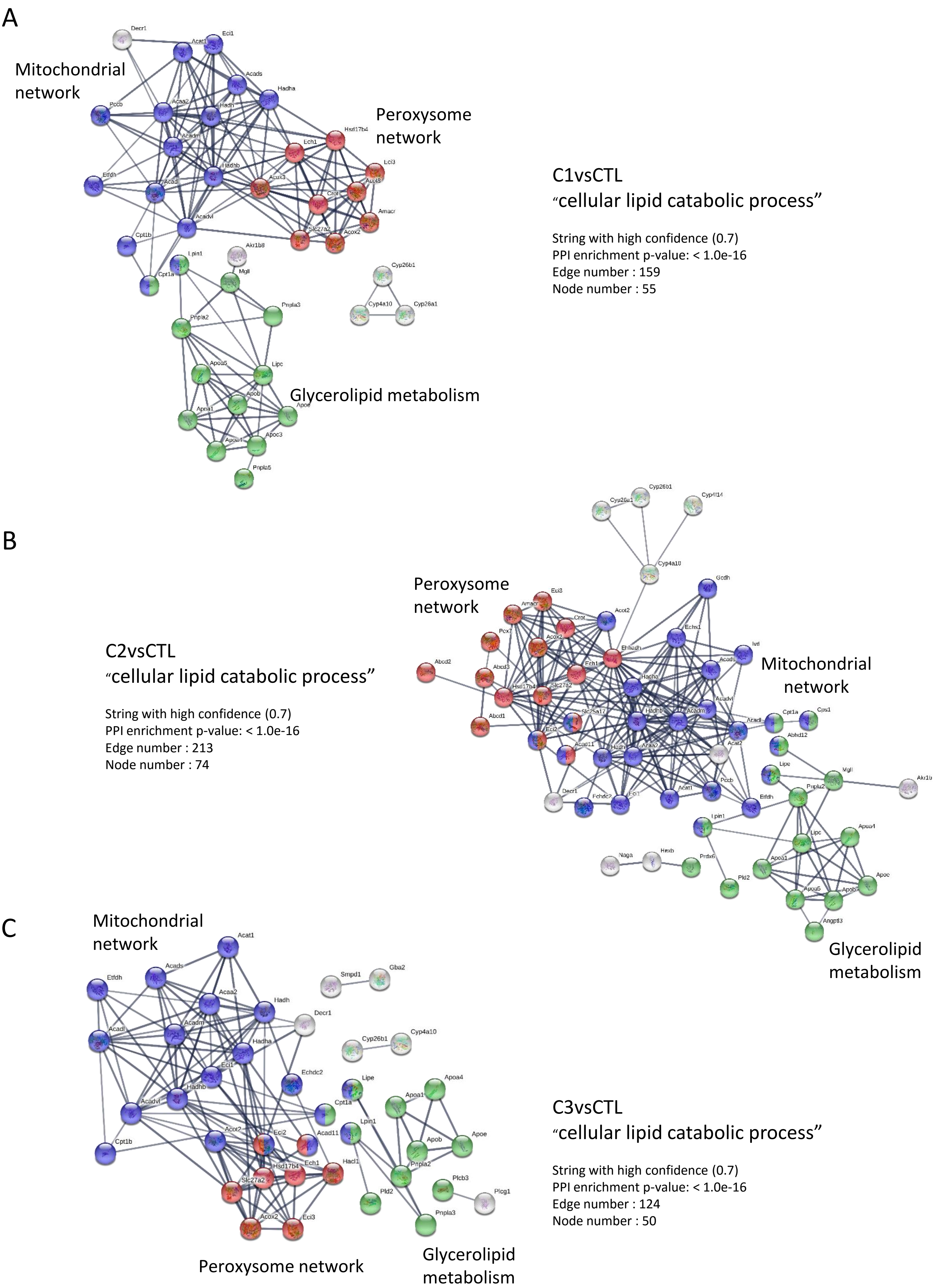
A
Mitochondrial network
Peroxysome network
Glycerolipid metabolism
C1vsCTL
"cellular lipid catabolic process"
String with high confidence (0.7)
PPI enrichment p-value: < 1.0e-16
Edge number : 159
Node number : 55
B
C2vsCTL
"cellular lipid catabolic process"
String with high confidence (0.7)
PPI enrichment p-value: < 1.0e-16
Edge number : 213
Node number : 74
Peroxysome network
Mitochondrial network
Glycerolipid metabolism
C
Mitochondrial network
Peroxysome network
Glycerolipid metabolism
C3vsCTL
"cellular lipid catabolic process"
String with high confidence (0.7)
PPI enrichment p-value: < 1.0e-16
Edge number : 124
Node number : 50

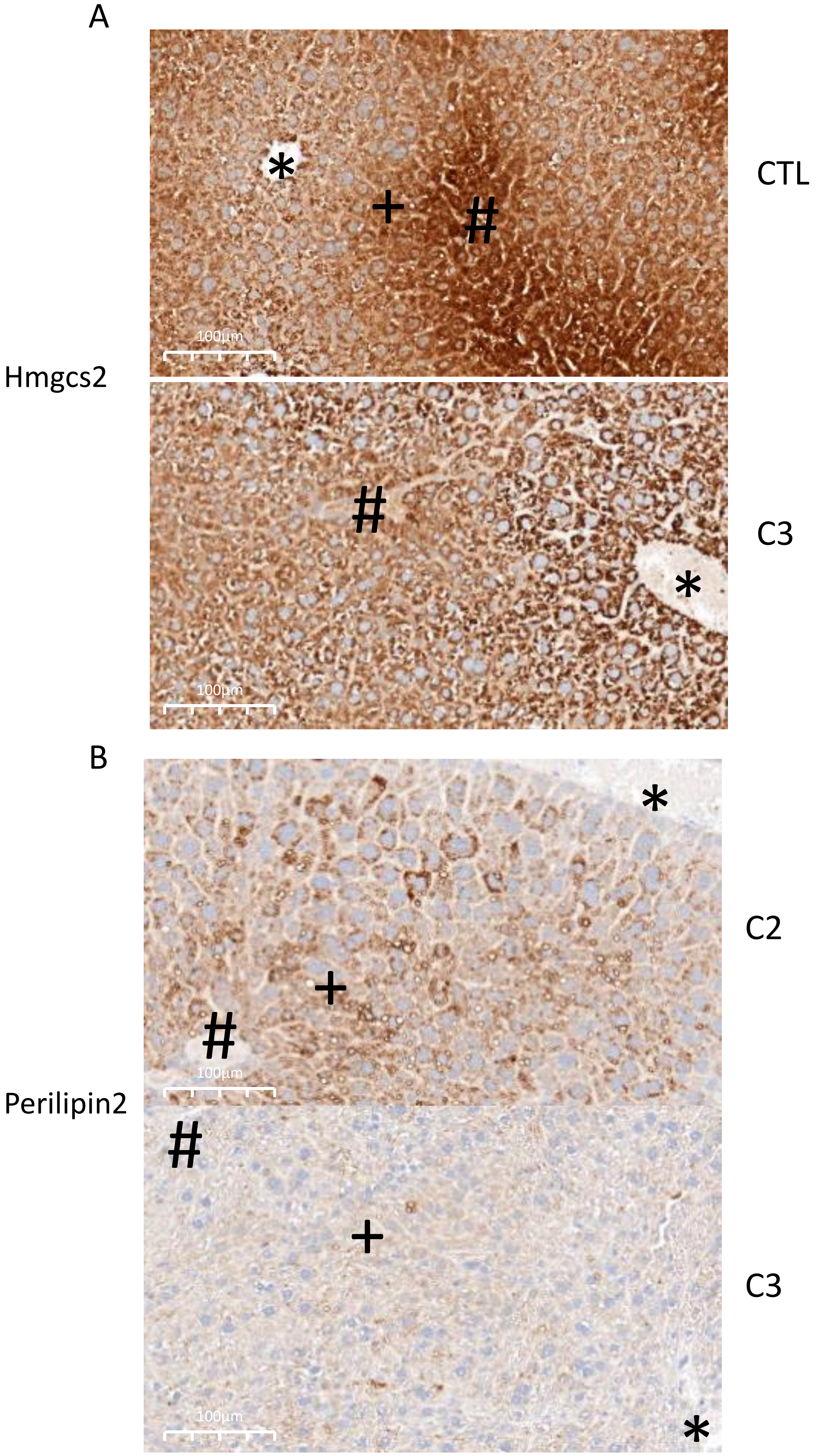
A
CTL
Hmgcs2
C3
B
C2
Perilipin2
C3
100µm
100µm
100µm
100µm
